# *Mtb* specific HLA-E restricted T cells are induced during *Mtb* infection but not after BCG administration in non-human primates and humans

**DOI:** 10.1101/2024.08.26.609630

**Authors:** Linda Voogd, Marjolein van Wolfswinkel, Iman Satti, Andrew D. White, Karin Dijkman, Anele Gela, Krista E. van Meijgaarden, Kees L.M.C. Franken, Julia L. Marshall, Tom H.M. Ottenhoff, Thomas J. Scriba, Helen McShane, Sally A. Sharpe, Frank A.W. Verreck, Simone A. Joosten

## Abstract

Novel vaccines targeting the world’s deadliest pathogen *Mycobacterium tuberculosis* (*Mtb*) are urgently needed as the efficacy of the Bacillus Calmette–Guérin (BCG) vaccine in its current use is limited. HLA-E is a virtually monomorphic unconventional antigen presentation molecule and HLA-E restricted *Mtb* specific CD8^+^ T cells can control intracellular *Mtb* growth, making HLA-E a promising vaccine target for *Mtb*. In this study, we evaluated the frequency and phenotype of HLA-E restricted *Mtb* specific CD4^+^/CD8^+^ T cells in the circulation and bronchoalveolar lavage fluid of two independent non-human primate (NHP) studies and from humans receiving BCG either intradermally or mucosally. BCG vaccination followed by *Mtb* challenge in NHPs did not affect the frequency of circulating and local HLA-E/*Mtb* CD4^+^ and CD8^+^ T cells, and we saw the same in humans receiving BCG. HLA-E/*Mtb* T cell frequencies were significantly increased after *Mtb* challenge in unvaccinated NHPs, which was correlated with higher TB pathology. Together, HLA-E/*Mtb* restricted T cells are minimally induced by BCG in humans and rhesus macaques (RMs) but can be elicited after *Mtb* infection in unvaccinated RMs. These results give new insights into targeting HLA-E as a potential immune mechanism against TB.

## 1. Introduction

Tuberculosis (TB) disease, caused by infection with *Mycobacterium tuberculosis* (*Mtb*), is a significant global health problem accounting for more than a million deaths each year[1]. Bacillus Calmette–Guérin (BCG) is the only licensed vaccine to protect against TB, however it has poor efficacy in adults in its current use. There is an urgent need for better protective vaccines against TB. Non-human primates (NHPs), specifically cynomolgus and rhesus macaques (CMs and RMs), are arguably the best pre-clinical models to evaluate novel vaccine strategies against TB and to increase our understanding of lung mucosal immune responses against *Mtb*. NHPs can develop the same pathology as human TB disease, including the formation of lung granulomas, lymph node involvement and the occurrence of latency[2,3].

Donor-unrestricted T cells (DURTs) recognize non-polymorphic antigen presentation molecules and most DURT subsets express invariant T cell receptors (TCRs)[4]. Vaccination approaches targeting DURTs can thus potentially induce protection irrespective of the genetic diversity in the human population. HLA-E restricted T cells belong to the DURT family because humans express two functional HLA-E alleles, HLA-E*01:01 and *01:03, that only differ in one amino acid located outside the peptide binding groove. This limited diversity suggests that both alleles have a comparable peptide binding repertoire[5,6]. HLA-E was first discovered as a ligand for the CD94/NKG2A(C) co-receptor inhibitory complex expressed on Natural Killer cells, which is an important surveillance mechanism to scan for, and subsequently clear, cells with defects in their antigen presentation machinery[7–9]. TCRs can also recognize HLA-E/peptide complexes[10] and HLA-E restricted CD8^+^ T cells have been identified in the circulation in patients with malignancies and infections, as we have shown previously in active TB (aTB) or *Mtb* infected (TBI) individuals with and without HIV co-infection[11–13]. Individuals with concomitant aTB and HIV infection demonstrated the highest frequency of circulating HLA-E restricted *Mtb* specific CD8^+^ T cells and revealed an unorthodox phenotype characterized by the secretion of T-helper 2 (Th2) associated cytokines, cytolysis of HLA-E/*Mtb* presenting target cells and inhibition of intracellular *Mtb* growth in *Mtb*-infected macrophages[13–22]. Moreover, in the absence of Qa-1^b^, the mouse equivalent of HLA-E, mice succumbed earlier from *Mtb* infection, suggesting a possible functional role of HLA-E in host defence[23].

The potential of HLA-E as a target for vaccination has been illustrated previously in RMs vaccinated with strain 68-1 rhesus cytomegalovirus (RhCMV) vectors encoding SIV antigens (called 68-1 RhCMV/SIV)[24]. Vaccination with this vector inhibited SIV replication and cleared SIV infection in more than half of the vaccinated RMs[25]. Importantly, the induction of MHC-E restricted CD8^+^ T cells following vaccination was essential to establish this protective effect[25,26]. Besides its virtual monomorphism, another advantage of targeting HLA-E by vaccination is that HLA-E is not downregulated upon HIV co-infection, in contrast to classical HLA-I molecules, which is an important benefit as infection with HIV and TB significantly overlap in endemic areas[27]. Together, these findings illustrate the potential for targeting HLA-E restricted T cells as a vaccination strategy against TB.

As individuals in TB endemic areas are routinely vaccinated with BCG, we sought to determine if HLA-E restricted *Mtb* specific T cells are induced in the circulation and bronchoalveolar lavage (BAL) fluid following BCG and/or *Mtb* challenge in two NHP studies and in humans after receiving BCG either intradermally (ID) or by aerosol. If induced, HLA-E might be a promising target to induce *de novo* T cell responses in BCG vaccinated individuals. Our results show that HLA-E/*Mtb* CD4^+^ and CD8^+^ T cell frequencies remained stable in the circulation of humans after receiving BCG, and in the BAL and circulation of BCG vaccinated and *Mtb* challenged RMs. Frequencies were increased in the BAL of unvaccinated and *Mtb* challenged RMs. These findings expand our knowledge on the induction of HLA-E restricted T cells after receiving BCG and upon *Mtb* infection in both the periphery as well as at the local site of infection.

## 2. Material and Methods

### 2.1 HLA-E TM folding and production

HLA-E*01:01 and *01:03 tetramers (TMs) were produced and correct folding of the monomers with the peptides was confirmed by staining LILRB1 expressing cells and mass spectrometry, as described previously[28]. Table 1 shows the HLA-E TMs used in each study.

**Table 1.**
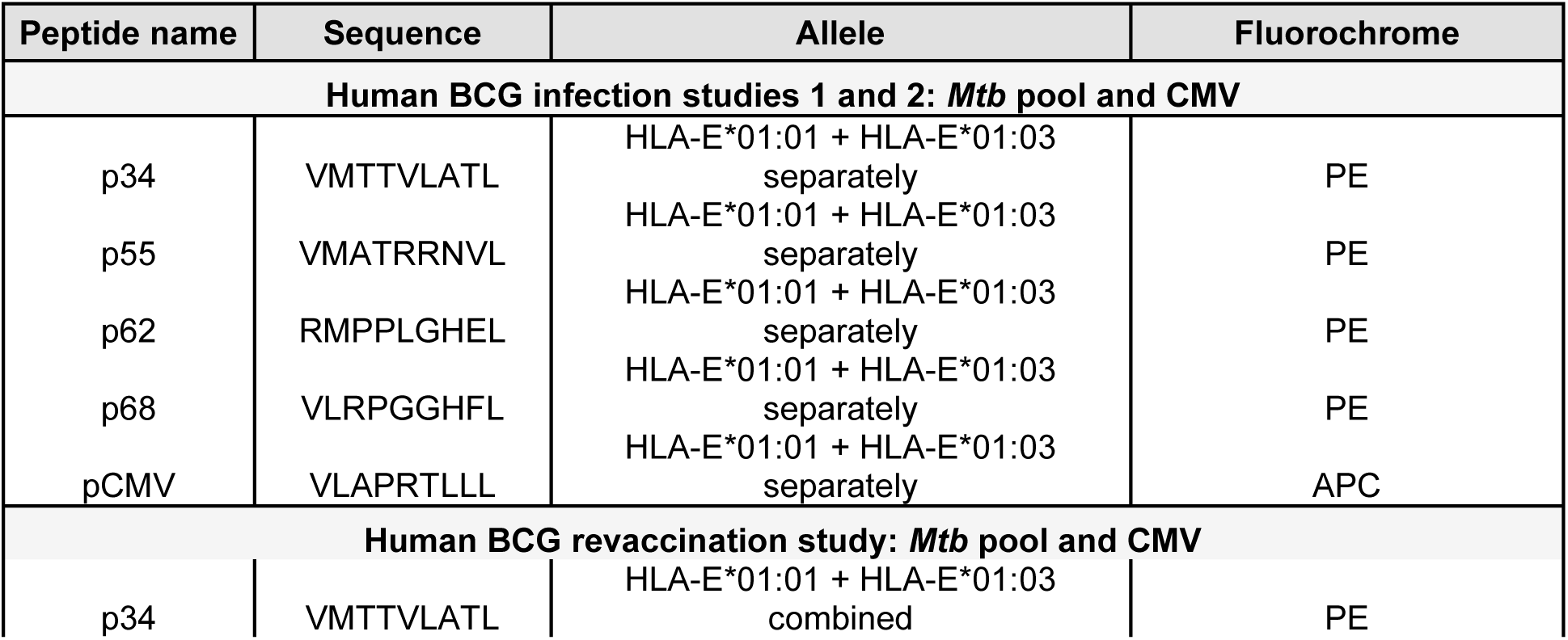
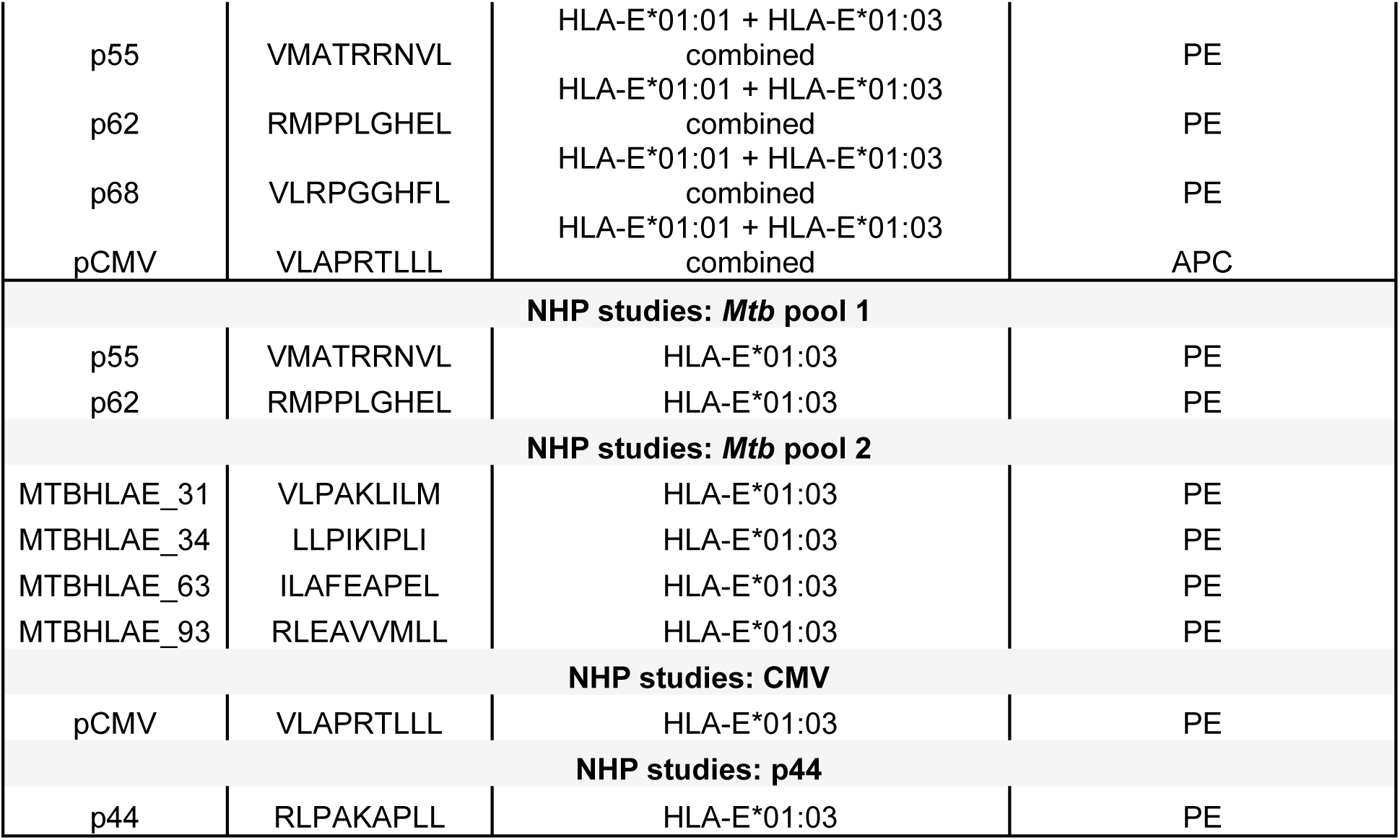
Overview of the HLA-E *Mtb* and CMV derived peptide (pools) used for HLA-E TM staining on PBMC and BAL samples in the human BCG revaccination and infection studies, RM studies 1 and 2 and the CM study.

### 2.2 Human BCG revaccination study

Eighty-two participants of a TBRU-supported BCG revaccination trial were enrolled at the South African Tuberculosis Vaccine Initiative, Cape Town, South Africa[29,30]. This trial was approved by the Medicines Control Council (MCC) of South Africa (now called South African Health Products Regulatory Authority, SAHPRA), by the University Hospital Cleveland Medical Center Institutional Board and the Human Research Ethics Committee of the University of Cape Town (387/2008). Written informed consent was obtained from all participants. The TBRU cohort comprised healthy tuberculin skin test positive, HIV uninfected adults who received routine BCG vaccination at birth. Participants were randomly assigned to two groups. Group 1 received at least six months of Isoniazid preventive therapy (IPT) before intradermal BCG revaccination and group 2 received IPT six months after intradermal BCG revaccination. Samples from twenty individuals of group 1 were used in the current study. Peripheral blood mononuclear cells (PBMCs) were collected and cryopreserved before BCG revaccination and 3, 5 and 52 weeks after BCG revaccination (Figure S1A for study overview). Cryopreserved PBMCs were thawed in pre-warmed (37°C) media containing DNAse (Sigma-Aldrich) and were washed in PBS. PBMCs were then stained with LIVE/DEAD^TM^ Fixable Near-IR Dead Cell Stain (Invitrogen) (1:1000 in PBS) for 20 min at room temperature (RT) in the dark. PBMCs were again washed in PBS and blocked with 12.5 µg/mL purified anti-CD94 antibody (BD Bioscience) followed by another wash in PBS/0.1% BSA. PBMCs were then stained with a pool of HLA-E*01:01 and *01:03 TMs (0.4 μg/TM) folded with the peptides shown in Table 1 for 15 min at 37°C in the dark. PBMCs were washed in PBS/0.1% BSA and were stained with APC-H7 CD14 (clone MOP9, BD Bioscience), APC-H7 CD19 (clone SJ25-C1, BD Bioscience), AlexaFluor700 CD3 (clone UCHT1, BioLegend), BV510 CD4 (clone RPA-T4, BioLegend), BV785 CD8 (clone SK1, BioLegend), BV605 CD26 (clone L272, BD Bioscience), BV650 CD161 (clone DX12, BD Bioscience), PE-Cy7 TRAV1.2 (clone 3C10, BioLegend), PE-Cy5 HLA-DR (clone L243, BioLegend), BV711 γδ TCR (clone B1, BD Bioscience), PE-CF594 CCR7 (clone 3D12, BD Bioscience) and PerCP-eFluor710 CD45RA (clone HI100, BioLegend) with pre-defined dilutions in BD Brilliant Stain Buffer (BD Bioscience) for 20 min at 4°C in the dark. PBMCs were washed in PBS/0.1% BSA, fixated in 1% paraformaldehyde, and acquired on a BD LSR-II flow cytometer. The gating strategy to determine HLA-E CD4^+^ and CD8^+^ T cell frequencies and their phenotype is shown in Figure S2A.

### 2.3 Controlled human BCG infection studies 1 and 2

Two controlled human BCG infection studies were performed at the University of Oxford (UK). The trial protocol of the first BCG controlled human infection study was approved by the Medicines and Healthcare products Regulatory Agency (EudraCT: 2015-004981-27) and the South-Central Oxford A Research Ethics Committee (REC) (15/SC/0716). Twelve BCG-naïve UK adults received the BCG Bulgaria strain (InterVax Ltd.) via aerosol inhalation of 1×10^7^ Colony Forming Units (CFU) using an Omron MicroAir mesh nebuliser and twelve adults via standard intradermal injection of 1×10^6^ CFU[31]. Blood was taken from all twenty-four volunteers at the start and seven and fourteen days after infection (Figure S1B for study overview). The trial protocol of the second aerosol BCG infection study was approved by the Central Oxford A REC (18/SC/0307) and is registered at ClinicalTrials.gov (NCT03912207). Six healthy BCG-naïve UK adults received the BCG Danish strain (AJVaccine) via aerosol inhalation of 1×10^7^ CFU using an Omron MicroAir mesh nebuliser. Bronchoalveolar lavage (BAL) and blood were taken fourteen days after receiving BCG from all individuals (Figure S1C for study overview). PBMCs were isolated and cryopreserved. Flow cytometric analysis was performed after thawing the PBMC samples whereas BAL samples were directly stained upon collection and isolation. BAL and PBMC (3×10^6^ cells each) were stained (1×10^6^ cells / well) with fixable LIVE/DEAD™ Vivid (Thermofisher) for 10 min at 40°C. Samples were then blocked with purified αCD94 (clone HP-3D9, BD Biosciences) and were stained with either: HLA-E*01:01, HLA-E *01:03 and CMV control TMs (Table 1) or left unstained to serve as gating control for TMs. Cells were incubated for 15 min at 37°C and were then washed before adding the surface staining antibody mix. The following antibodies were added to all cells: CD3-AF700 (Clone UCHT1, Thermofisher), CD4-FITC (Clone: RPA-T4, BioLegend), CD8-APC/H7 (Clone SK1, BD Bioscience), CD14-Pacific blue (Thermofisher) and CD19-Pacific blue (Thermofisher). Cells were incubated with the surface antibody mix for 30 min at 4°C and were then washed before being acquired on LSR Fortessa v.2 Std X20 flow cytometer using BD FACSDiva 8.0.2 and the data was analysed on FlowJo v9. The gating strategy to determine HLA-E CD4^+^ and CD8^+^ T cell frequencies in human PBMC and BAL for studies 1 and 2 is shown in Figure S2B.

### 2.4 Rhesus macaque study 1: Ethics, animal care, vaccination and pathology assessment

Archived samples from the study published by White *et al.* were used in the current study[32]. Study design and animal welfare procedures were approved by the UK Health Security Agency Animal Welfare and Ethical Review Body and was authorised under UK Home Office Licence P219D8D1A. Nineteen healthy Indian-type rhesus macaques (*Macaca mulatta*) from a closed UK breeding colony were housed in groups in accordance with the UK Home Office Code of Practice and NC3Rs guidelines on Primate Accommodation. Six animals received the BCG Danish strain 1331 via intradermal injection into the upper left arm and six animals received BCG by aerosol exposure using an Omron MicroAir mesh nebuliser with an estimated dose of 2-8×10^6^ CFU/mL. The control group consisted of seven unvaccinated animals. Twenty-one weeks after BCG vaccination, all animals were challenged with a calculated single ultra-low dose of 3 CFU *Mtb* Erdman K01 via aerosol delivery. The apparatus and procedure for aerosol delivery of *Mtb*, including the calculation of the presented dose were performed as described previously[32]. Blood from all RMs was collected and PBMCs were cryopreserved. Samples at the time points shown in Figure S1D were shipped to the LUMC, The Netherlands for flow cytometry analysis. PBMCs were thawed for staining with HLA-E*01:03 TMs folded with *Mtb* pool 1 and pool 2, p44 and CMV (Table 1) and cell surface markers as outlined in detail below. Samples were acquired on a BD FACS Lyric 3L12C (BD Biosciences).

### 2.5 Rhesus macaque study 2: Ethics, animal care, vaccination, and pathology assessment

Archived samples from the study published by Dijkman *et al.* were used in the current study[33]. Ethical approval of the study protocol was obtained from the independent ethics committee Dier Experimenten Commissie (DEC) (761subB) and the institutional animal welfare body of the Biomedical Primate Research Center (BPRC). The approved housing and animal care procedures are described previously[33]. Twenty-four healthy male Indian-type rhesus macaques (*Macaca mulatta*) from the in-house breeding colony were stratified into three groups of eight animals. BCG vaccination or placebo was randomly assigned to each group. Two of the three groups received between 1.5-6.0×10^5^ CFU of the BCG strain Sofia (InterVax Ltd.), either via intradermal injection or via endobronchial instillation (referred to as mucosal vaccination) into the lower left lung lobe. Animals receiving saline via endobronchial instillation served as the control group. Twelve weeks after BCG vaccination or saline treatment all animals received a weekly calculated limiting dose of 1 CFU *Mtb* Erdman K01 strain for eight consecutive weeks. Infection was confirmed with IFN-γ ELISpot. Blood and BAL samples were collected, PBMCs and BAL cells were isolated and cryopreserved. Samples at the time points shown in Figure S1E were shipped to the LUMC, The Netherlands. PBMCs and BAL cell samples were thawed for staining with HLA-E*01:03 TMs folded with *Mtb* pool 1 and pool 2, p44 and CMV (Table 1) and cell surface markers as outlined in detail below. Samples were acquired on a BD FACS Lyric 3L12C (BD Biosciences). Bacterial loads (i.e., CFU counts) in lung tissues and post-mortem pathology scores (i.e., lesion size and granuloma formation) were determined at the BPRC.

### 2.6 Cynomolgus macaque study: Ethics, animal care, vaccination and pathology assessment

Archived samples from the study published by White *et al.* were used in the current study[34]. Study design and animal housing was approved by the Establishment Animal Welfare and Ethical Review Committee and authorised under UK Home Office Project License P219D8D1A. The approved housing and animal care procedures are described previously[34]. Nine cynomolgus macaques (*Macaca fascicularis*) from a closed UK breeding colony were stratified into three groups of three animals. One group received between 2-8×10^5^ CFU/mL BCG Danish strain 1331 via intradermal injection into the upper left arm. Twenty-one weeks after vaccination all animals were challenged with a calculated dose of 5 CFU *Mtb* Erdman strain K01 via aerosol delivery. Sixteen weeks after *Mtb* challenge, all groups were challenged with 106-107 (TCID)_50_ SIVmac32H 11/88. Infection with SIV was confirmed via culturing PBMCs with C8166 cells to examine the cytopathic effect. Blood from all animals was collected, and PBMCs were isolated and cryopreserved at the time points shown in Figure S1F. Cryopreserved PBMCs were shipped to the LUMC, The Netherlands for flow cytometry analysis. PBMCs were thawed for staining with HLA-E*01:03 TMs folded with *Mtb* pool 1 and 2, p44 and CMV (Table 1) and cell surface markers as outlined in detail below. Samples were acquired on a BD FACS Lyric 3L12C (BD Biosciences).

### 2.7 Flow cytometry staining, acquisition and analysis of NHP PBMC and BAL samples

NHP samples were processed at the Biosafety Level 3 (BSL3) laboratory at the LUMC, The Netherlands. PBMC and BAL samples were thawed and rested for 2 hrs in PBS with 0.2 mg/mL DNase I (Roche Diagnostics GmbH) at 37°C with 5% CO_2_ to adhere monocytic cells. Cells were washed (10 min, 450 x g) and stained using previously developed and optimized staining procedures for high dimensional flow cytometry panels[35]. Samples were stained with LIVE/DEAD™ Fixable Violet (Invitrogen) according to manufacturer’s instructions in PBS for 30 min at room temperature (RT) in the dark. Cells were then washed (5 min, 450 x g) with PBS, blocked with 5% pooled normal human serum in PBS for 10 min at RT to block Fc receptors and prevent non-specific binding and washed once with PBS/0.1% BSA. Cells were subsequently stained with an antibody cocktail containing True-Stain Monocyte Blocker (1:20, BioLegend) to block non-specific binding of tandem dyes to monocytes, Brilliant Stain Buffer Plus (1:10, BD Biosciences) and the chemokine receptor antibodies PE-Cy7 CCR4 (1:100, clone 1G1, BD Biosciences), BV605 CCR6 (1:100, clone 11A9, BD Biosciences), BV785 CCR7 (1:50, clone G043H7, BioLegend) and APC-Cy7 CXCR3 (1:50, clone G025H7, BioLegend) in PBS/0.1% BSA for 30 min at 37°C in the dark. After incubation, cells were washed once with PBS/0.1% BSA and incubated with the HLA-E*01:03 TM conditions (5.4 µg/mL per TM) shown in Table 1 in PBS/0.1% BSA for 30 min at 37°C in the dark. Peptide p44 is a high affinity *Mtb*-derived HLA-E restricted peptide that adopts a similar conformation in the peptide binding groove as classical HLA-I leader sequence derived peptides[36] and served as a control. Cells were then washed once with PBS/0.1% BSA, fixated with 1% paraformaldehyde (Pharmacy LUMC, Leiden) for 10 min at RT and washed with PBS/0.1% BSA. Cells were subsequently stained with an antibody cocktail containing Brilliant Stain Buffer Plus (1:10) and the surface antibodies BV421 CD14 (1:100, clone M5E2, BD Biosciences) and BV421 CD19 (1:200, clone 2H7, BD Biosciences) as dump channel together with the LIVE/DEAD stain, PerCP-Cy5.5 CD3 (1:25, clone SP34-2, BD Biosciences), R718 CD4 (1:50, clone L200, BD Biosciences), FITC CD8 (1:50, clone SK1, BD Biosciences), BV510 CD45RA (1:200, clone 5H9, BD Biosciences), BV711 CD16 (1:100, clone 3G8, BD Biosciences) and APC NKG2A (1:100, clone Z199, Beckman Coulter) in PBS/0.1% BSA for 30 min at 4°C in the dark. For the BAL samples, because of high autofluorescence, BV605 CD8 (1:100, clone SK1, BD Biosciences) was used instead of FITC CD8 and therefore CCR6 was not included in the BAL sample analysis. After incubation, cells were washed twice with PBS/0.1% BSA, fixated with 1% paraformaldehyde for 10 min at RT and washed twice with PBS/0.1% BSA. Cells were then resuspended in PBS/0.1% BSA and acquired on a BD FACS Lyric 3L12C (BD Biosciences). Flow cytometry data was analysed using FlowJo v10.9.0. Subsequent (statistical) analysis was performed in GraphPad Prism v9.3.1. The gating strategy to determine HLA-E CD4^+^ and CD8^+^ T cell frequencies and their phenotype in PBMC and BAL samples are shown in Figure S3A and B, respectively.

## 3 Results

### 3.1 The frequency of circulating HLA-E/Mtb CD4^+^ and CD8^+^ T cells remains stable in humans and RMs after receiving BCG

The frequency of circulating HLA-E/*Mtb* CD4^+^ and CD8^+^ T cells remained stable after intradermal BCG revaccination in healthy adults (Figure 1A), similar to the stability observed in BCG naïve humans receiving primary intradermal or aerosol BCG (Figure 1B). BCG administered as a boost or prime therefore did not induce HLA-E/*Mtb* CD4^+^ and CD8^+^ T cells in humans.

**Figure 1.**
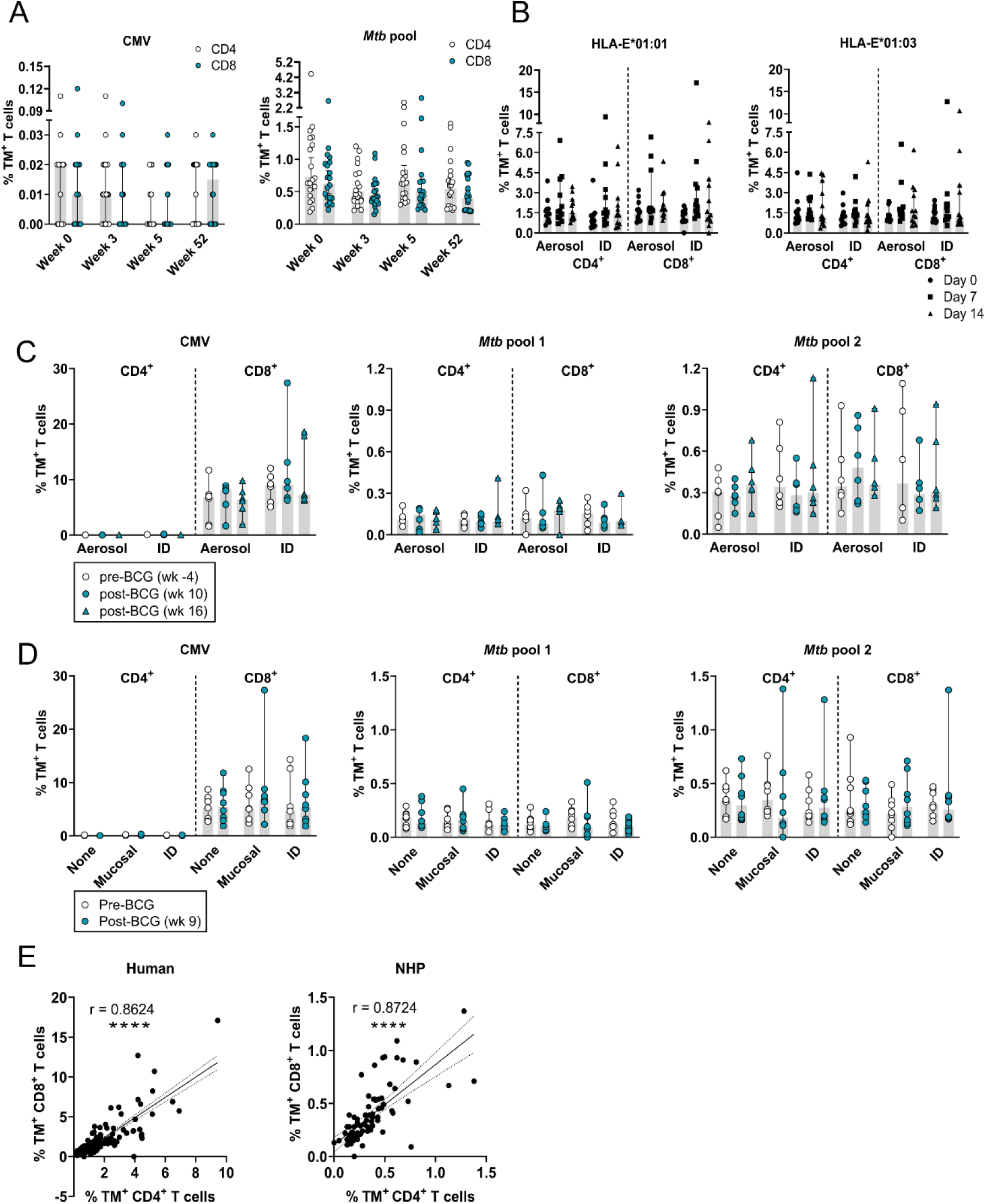
HLA-E/*Mtb* and CMV CD4^+^ and CD8^+^ T cell frequencies in the circulation remain stable after receiving BCG in humans and RMs. Peptide pools for HLA-E TM staining on human and RM samples are shown in Table 1. **(A)** HLA-E*01:03 and *01:01 CD4^+^ and CD8^+^ T cell frequencies for CMV (left) and the *Mtb* pool (right) at week 0 and 3, 5 and 52 weeks after intradermal BCG revaccination in healthy TST^+^ HIV^-^ volunteers (n=20); **(B)** HLA-E*01:01 (left) and *01:03 (right) CD4^+^ and CD8^+^ T cell frequencies for the *Mtb* pool on day 0 (circles) and 7 (squares) and 14 (triangles) days after aerosol BCG inhalation (n=12) or intradermal BCG administration (n=12) in healthy volunteers; **(C)** HLA-E*01:03 CD4^+^ and CD8^+^ T cell frequencies for CMV (left), *Mtb* pool 1 (middle) and 2 (right) in RM study 1 at the pre-vaccination time point (white circles), 10 (blue circles) and 16 weeks post-vaccination (blue triangles); **(D)** Same as C, but then for RM study 2 at the pre-vaccination time point (white circles) and 9 weeks post-vaccination (blue circles); **(E)** Correlating HLA-E CD4^+^ T cell frequencies (X-axis) and HLA-E CD8^+^ T cell frequencies (Y-axis) at all time points in the human studies for the *Mtb* pool (left) and in RM study 1 and 2 combined for *Mtb* pool 2 (right). Shaded bars represent the median frequency, and the error bars represent the 95% confidence interval. Significance was tested using a repeated measures (RM) two-way ANOVA with multiple comparison correction (A-D) and a Spearman’s rank correlation (E). * = *p*<0.05, ** = *p*<0.01, **** = *p*<0.0001.

Representative density plots for p44 (negative control), CMV and *Mtb* pool 1 and 2 before and after BCG vaccination in RMs are shown in Figure S4A and B, respectively. The frequency of circulating HLA-E/*Mtb* CD4^+^ and CD8^+^ T cells was unchanged post-BCG vaccination compared to pre-vaccination in both RM study 1 and 2 with no differences observed between administration routes (Figure 1C and D). HLA-E CMV TMs were only recognized by CD8^+^ T cells, whereas HLA-E/*Mtb* TMs were recognized by both CD4^+^ and CD8^+^ T cells in RMs and the recognition between CD4^+^ and CD8^+^ HLA-E/*Mtb* T cells was strongly correlated in both humans and RMs (Figure 1E). The frequency of HLA-E/CMV CD8^+^ T cells in the circulation was overall higher compared to HLA-E/*Mtb* CD4^+^ and CD8^+^ T cells (10% vs. 0.3%). Combined, BCG administration in RMs did not increase the frequency of HLA-E/*Mtb* T cells in the circulation irrespective of the administration route.

### 3.2 The frequency of circulating HLA-E/Mtb CD4^+^ and CD8^+^ T cells does not change after Mtb challenge in unvaccinated and BCG vaccinated RMs

Representative density plots for *Mtb* pool 2 (which shares the same recognition profile as *Mtb* pool 1; see Figure S5 for results on *Mtb* pool 1) on pre-and post-*Mtb* challenge samples from one aerosol BCG vaccinated RM are shown in Figure S4C. *Mtb* challenge with or without prior BCG vaccination did not affect the frequency of circulating HLA-E/*Mtb* CD4^+^ and CD8^+^ T cells in both RM studies and no differences were observed between the BCG administration routes (Figure 2A and B). As such, the frequency of circulating HLA-E/*Mtb* CD4^+^ and CD8^+^ T cells was not changed after *Mtb* challenge only or following BCG vaccination in RMs.

**Figure 2.**
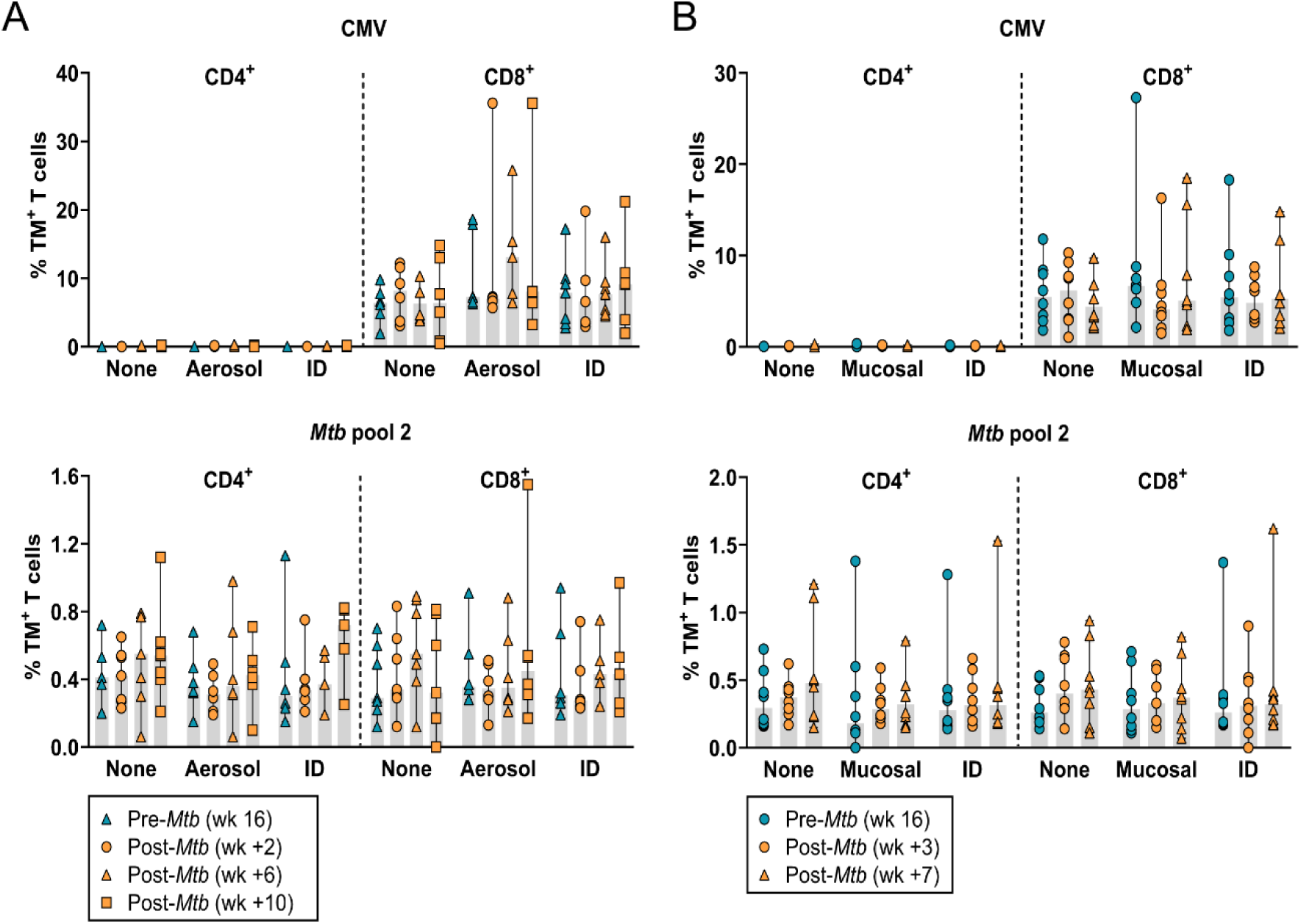
*Mtb* challenge only or following BCG vaccination in RMs results in stable HLA-E/*Mtb* and CMV CD4^+^ and CD8^+^ T cell frequencies in the circulation. Peptide pools for HLA-E TM staining on RM samples are shown in Table 1. **(A)** HLA-E*01:03 CD4^+^ and CD8^+^ T cell frequencies for CMV (upper panel) and *Mtb* pool 2 (lower panel) in RM study 1 at the pre-challenge time point (blue triangles) and 2 (orange circles), 6 (orange triangles) and 10 weeks (orange squares) post-*Mtb* challenge; **(B)** Same as A, but then in RM study 2 at the pre-challenge time point (blue circles) and 3 (orange circles) and 7 weeks (orange triangles) post-*Mtb* challenge. Shaded bars represent the median frequency, and the error bars represent the 95% confidence interval. Significance was tested using a repeated measures (RM) two-way ANOVA with multiple comparison correction. * = *p*<0.05, ** = *p*<0.01, **** = *p*<0.0001.

### 3.3 Increased frequency of HLA-E/Mtb CD4^+^ and CD8^+^ T cells in the BAL after Mtb challenge in unvaccinated RMs

Representative density plots for p44, CMV, *Mtb* pool 1 and 2 in the BAL from an unvaccinated RM after *Mtb* challenge are shown in Figure 3A. Both in RMs and humans, the frequency of HLA-E/*Mtb* CD4^+^ and CD8^+^ T cells was significantly higher in the BAL compared to the circulation, including for HLA-E/CMV CD8^+^ T cells in RMs (Figure 3B and C). Similar as in the circulation, the frequency of HLA-E/CMV CD8^+^ T cells was higher than HLA-E/*Mtb* CD4^+^ and CD8^+^ T cells in the BAL (Figure 3B). In contrast to BCG vaccination, *Mtb* challenge markedly and significantly increased the frequency of HLA-E/*Mtb* CD4^+^ and CD8^+^ T cells in the BAL of unvaccinated RMs (Figure 3D). There was a positive and significant correlation between the PA scores and CFU counts in the lung, primary and secondary lobe of all RMs, which was most pronounced in unvaccinated RMs (Figure 3E). HLA-E/*Mtb* CD3^+^ T cell frequencies in the BAL and the PA scores in the total lung and primary lobe were significantly correlated as well (Figure 3F), which suggests that BCG vaccination can contribute to protection, reflected by the lower PA scores in the vaccinated RMs, but likely by other mechanism than HLA-E, if increased frequencies of HLA-E/*Mtb* restricted T cells is required for protection.

**Figure 3.**
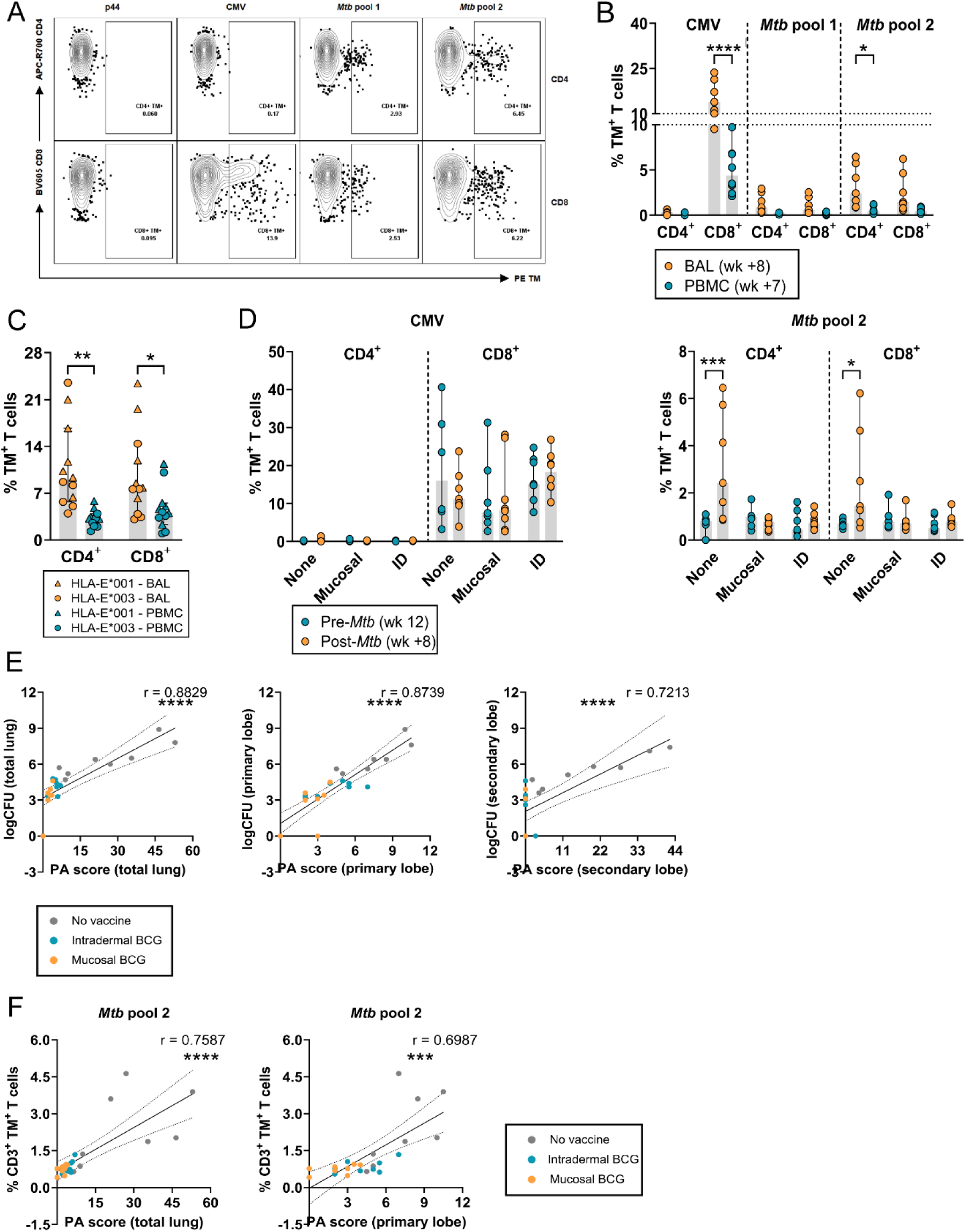
HLA-E/*Mtb* and CMV CD4^+^ and CD8^+^ T cell frequencies are increased after *Mtb* challenge in BAL of BCG unvaccinated RMs. Peptide pools for HLA-E TM staining on RM and human samples are shown in Table 1. **(A)** Density plots for HLA-E p44, CMV and *Mtb* pool 1 and 2 CD4^+^ and CD8^+^ T cell frequencies in a BAL sample of one representative unvaccinated RM of study 2 post-*Mtb* challenge. HLA-E TMs in PE are shown on the X-axis and CD4 and CD8 are shown on the Y-axis; **(B)** HLA-E*01:03 CD4^+^ and CD8^+^ T cell frequencies for CMV and *Mtb* pool 1 and 2 in BAL (orange) and PBMCs (blue), 8 and 7 weeks, respectively, post-*Mtb* challenge in unvaccinated RMs of RM study 2 (n=8); **(C)** HLA-E*01:01 (triangles) and *01:03 (circles) CD4^+^ and CD8^+^ T cell frequencies in BAL (orange) and in PBMCs (blue) for the *Mtb* pool, 14 days after aerosol BCG inhalation in healthy volunteers (n=12); **(D)** HLA-E*01:03 CD4^+^ and CD8^+^ T cell frequencies in BAL for CMV (left) and *Mtb* pool 2 (right) in RM study 2, 12 weeks post-BCG vaccination (blue) and 8 weeks post-*Mtb* challenge (orange); **(E)** Correlating the CFU counts (Y-axis) and the PA scores (X-axis) in the total lung (left), primary lobe (middle) and secondary lobe (right) for RM study 2. Grey dots represent the unvaccinated group, blue dots the intradermal BCG vaccinated group, and orange dots the mucosal BCG vaccinated group. Dotted lines represent the 95% confidence interval; **(F)** Same as E, but then correlating HLA-E*01:03 *Mtb* CD3^+^ T cell frequencies in BAL (Y-axis) and the PA scores in the total lung (left) and primary lobe (right) (X-axis) 8 weeks post-*Mtb* challenge. Shaded bars represent the median frequency, and the error bars represent the 95% confidence interval. Significance was tested using a repeated measures (RM) two-way ANOVA with multiple comparison correction (A-D) and a Spearman’s rank correlation (E-F). * = *p*<0.05, ** = *p*<0.01, **** = *p*<0.0001.

### 3.4 HLA-E/Mtb CD3^+^ T cells have a similar memory phenotype but altered chemokine receptor expression compared to total T cells

Circulating HLA-E/*Mtb* CD4^+^ and CD8^+^ T cells had a similar distribution of memory subsets compared to total T cells in both RM studies, which was not changed upon BCG vaccination or *Mtb* challenge (Figure 4A and B). In contrast, HLA-E/CMV CD8^+^ T cells had a dominant effector memory (EM) and terminally differentiated (EMRA) phenotype that did not change after BCG vaccination and *Mtb* challenge in both RM studies (Figure 4A and B). Both CD4^+^ and CD8^+^ T cells can recognize HLA-E/*Mtb* (Figure 1E), therefore CCR expression was evaluated on CD3^+^ T cells. HLA-E/*Mtb* T cells had a significantly higher expression of CCR4 and lower expression of CXCR3 compared to HLA-E/CMV T cells and a higher expression of CXCR3 compared to total T cells post-BCG vaccination (RM study 1 in Figure 4C and RM study 2 in 4D). The expression of CCR6 was similar on HLA-E/CMV, HLA-E/*Mtb* T cells and total T cells in RM study 1 (Figure 4C), but significantly increased on HLA-E/CMV T cells compared to HLA-E/*Mtb* T cells in RM study 2 (Figure 4D). Similar findings for CCR expression were found in the BAL, although the overall expression of CXCR3 was higher in the BAL compared to the circulation (Figure 4E). Whereas the expression of CCR4, CCR6 and CXCR3 on HLA-E/*Mtb* T cells in the circulation and the BAL did not change upon *Mtb* challenge in unvaccinated and vaccinated RMs (Figure 4F, G and H), HLA-E/CMV T cells in unvaccinated RMs in study 2 had decreased expression of CXCR3 and CCR6 in the circulation and of CXCR3 in the BAL after *Mtb* challenge compared to vaccinated RMs (Figure 4G and H). These findings reveal that (i) BCG vaccination followed by *Mtb* challenge did not change the memory phenotype of HLA-E/*Mtb* T cells in the circulation and that (ii) HLA-E/*Mtb* T cells have different CCR expression levels compared to HLA-E/CMV T cells and total T cells, suggesting pathogen specific expression.

**Figure 4.**
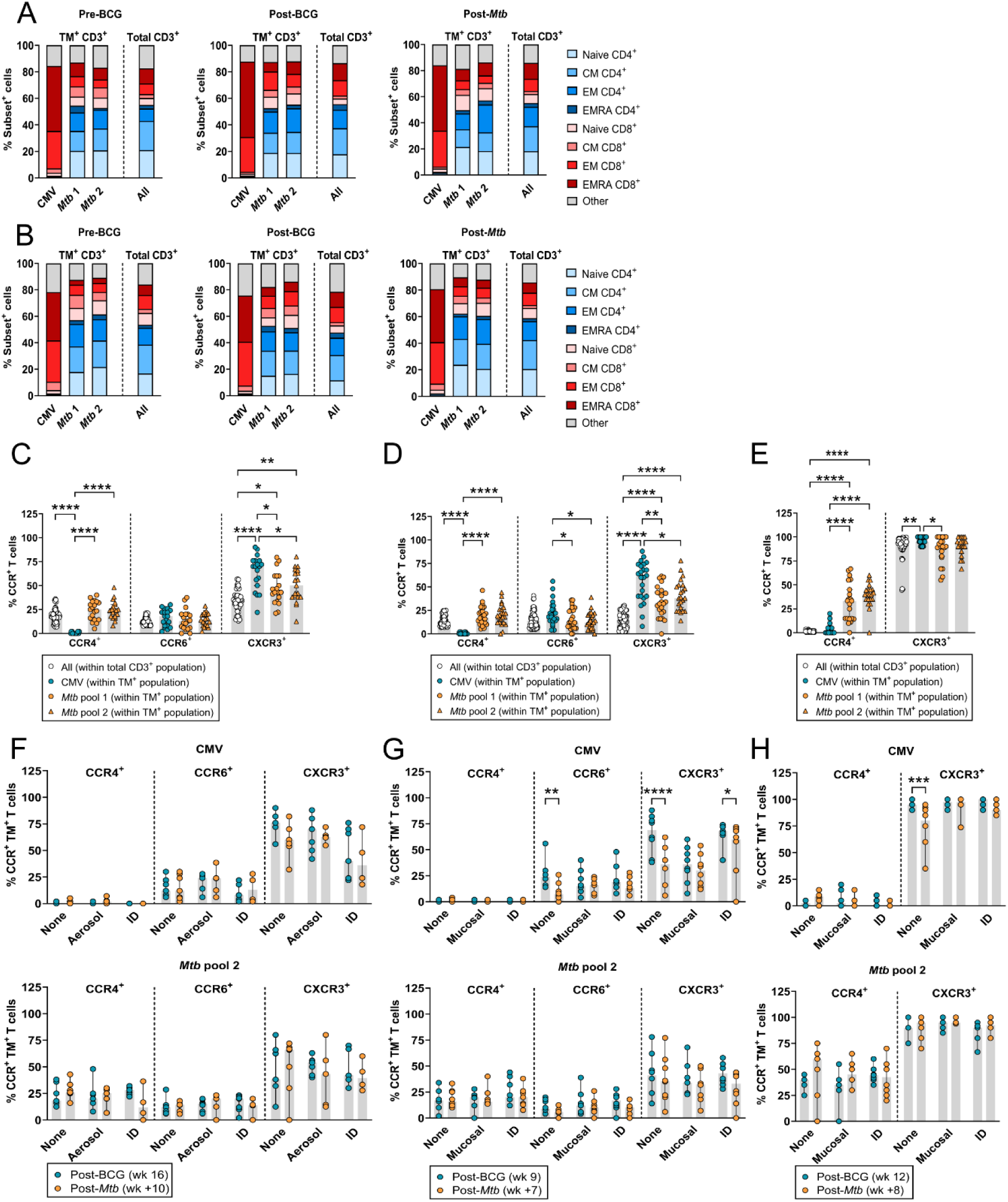
Phenotypic analysis of HLA-E/Mtb CD4^+^ and CD8^+^ T cells after BCG vaccination and *Mtb* challenge in RMs. Peptide pools for HLA-E TM staining on RM samples are shown in Table 1. **(A)** Memory subset identification of HLA-E*01:03 *Mtb* and CMV CD3^+^ T cells relative to total CD3^+^ T cells in the circulation of RM study 1 (n=19), 4 weeks pre-BCG vaccination (left), 16 weeks post-BCG vaccination (middle) and 10 weeks post-*Mtb* challenge (right); **(B)** Same as A, but then in RM study 2 (n=24) 2 weeks pre-BCG vaccination (left), 9 weeks post-BCG vaccination (middle) and 7 weeks post-*Mtb* challenge (right); **(C)** CCR4, CCR6 and CXCR3 expression on HLA-E*01:03 CMV (blue circles), *Mtb* pool 1 (orange circles) and 2 (orange triangles) CD3^+^ T cells relative to total CD3^+^ T cells (white circles) in the circulation 16 weeks post-BCG vaccination in RM study 1 (n=19); **(D)** Same as C, but then for RM study 2 (n=24) 9 weeks post-BCG vaccination; **(E)** Same as D (without CCR6), but then in the BAL 12 weeks post-BCG vaccination; **(F)** CCR4, CCR6 and CXCR3 expression on HLA-E*01:03 CMV (upper panel) and *Mtb* pool 2 (lower panel) CD3^+^ T cells in the circulation 16 weeks post-BCG vaccination (blue) and 10 weeks post-*Mtb* challenge (orange) in RM study 1; **(G)** Same as F, but then for RM study 2, 9 weeks post-BCG vaccination (blue) and 7 weeks post-*Mtb* challenge (orange); **(H)** Same as G, but then in the BAL 12 weeks post-BCG vaccination (blue) and 8 weeks post-*Mtb* challenge (orange). Shaded bars and stacked bars represent the median and mean frequency, respectively, and the error bars represent the 95% confidence interval. Significance was tested using a repeated measures (RM) two-way ANOVA with multiple comparison correction (C – H). * = *p*<0.05, ** = *p*<0.01 *** = *p*<0.001, **** = *p*<0.0001.

### 3.5 Simian Immunodeficiency Virus (SIV) co-infection in Mtb-challenged cynomolgus macaques and HLA-E/Mtb CD4^+^ and CD8^+^ T cell frequencies

Although only three CMs per group, BCG vaccinated, *Mtb* and SIV co-infected CMs tended to have a lower frequency of HLA-E/*Mtb* CD4^+^ and CD8^+^ T cells compared to unvaccinated *Mtb* and SIV co-infected CMs, both after *Mtb* and SIV infection (Figure 5A). This was not observed for HLA-E/CMV CD8^+^ T cells suggesting some level of protection by BCG. HLA-E/*Mtb* and CMV CD4^+^ and CD8^+^ T cell frequencies were similar in *Mtb* SIV co-infected and *Mtb* only infected CMs, both in the absence of BCG vaccination, suggesting no effect of SIV co-infection (Figure 5B), and after *Mtb* challenge before SIV infection (Figure 5C), similar to the findings in RMs (Figure 2).

**Figure 5.**
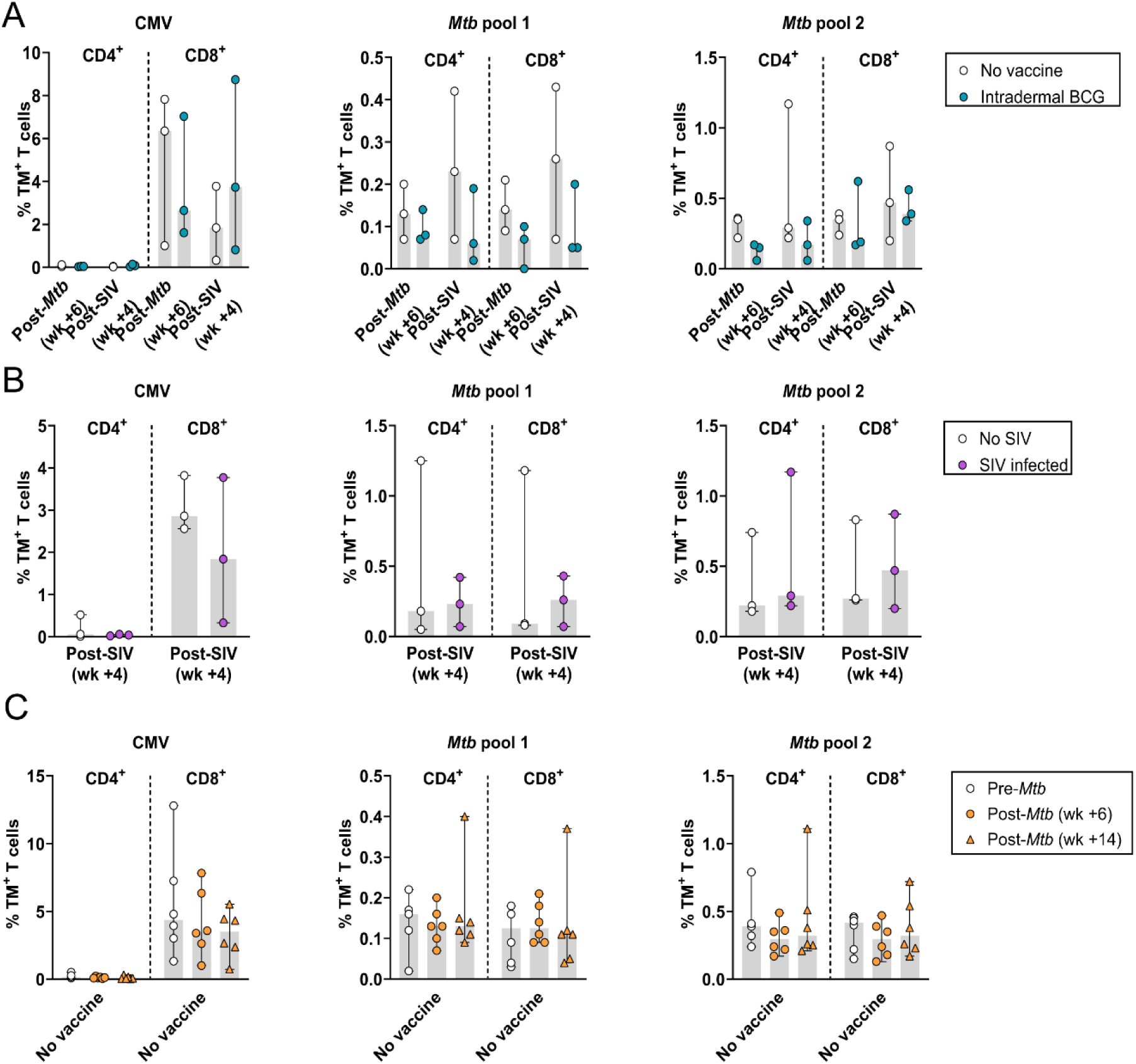
Effect of *Mtb*/SIV co-infection on HLA-E/*Mtb* and CMV CD4^+^ and CD8^+^ T cell frequencies in cynomolgus macaques (CMs). Peptide pools for HLA-E TM staining on CM samples are shown in Table 1. 3 CMs were vaccinated with BCG and were challenged with *Mtb* followed by SIV infection, 3 CMs were challenged with *Mtb* followed by SIV infection without BCG vaccination and 3 CMs were challenge with *Mtb* only. **(A)** HLA-E*01:03 CMV (left), *Mtb* pool 1 (middle) and 2 (right) CD4^+^ and CD8^+^ T cell frequencies 6 weeks post-*Mtb* challenge and 4 weeks post-SIV challenge in unvaccinated (white) and BCG vaccinated (blue) CMs; **(B)** Same as A, but then 4 weeks post-SIV challenge in unvaccinated and *Mtb* challenged CMs, either without (white) or with (purple) subsequent SIV infection; **(C)** Same as A, but then pre-*Mtb* challenge (white circles) and 6 (orange circles) and 14 weeks (orange triangles) post-*Mtb* challenge (before SIV infection) in six unvaccinated CMs. Shaded bars represent the median frequency, and the error bars represent the 95% confidence interval. Significance was tested using a repeated measures (RM) two-way ANOVA with multiple comparison correction.

## 4 Discussion

We determined HLA-E restricted *Mtb* specific T cell frequencies and their memory phenotypes after BCG vaccination and/or *Mtb* infection in NHPs and in harmonized controlled BCG infection studies in humans. Mucosal and intradermal BCG vaccination did not modulate the frequency of circulating and local HLA-E/*Mtb* CD4^+^ and CD8^+^ T cells in RMs, nor in humans receiving aerosol or intradermal BCG. Previously, intradermal BCG revaccination was found to have no effect on the frequencies of various DURT subsets in the circulation of adults, including MAIT cells, γδ T cells, NKT cells, CD1b and germline-encoded mycolyl-reactive (GEM) T cells[29]. However, γδ T cells were increased after primary vaccination in infants[29]. Our findings show that, similar to other DURT subsets, frequencies of HLA-E/*Mtb* restricted T cells were unchanged after BCG revaccination. A previous study showed that intravenous BCG vaccination in RMs induced the highest level of *Mtb* specific T cell responses in the circulation and the BAL and induced the lowest CFU counts compared to intradermal and aerosol. Intravenous BCG might also be more efficient at inducing HLA-E/*Mtb* T cells, though this has to be evaluated in future studies[37]. As the aligned studies in RMs and humans show an almost identical effect of BCG on HLA-E/*Mtb* T cell frequencies, this underscores the relevance of RMs as a model for human mycobacterial infections.

*Mtb* challenge significantly increased frequencies of HLA-E/*Mtb* CD3^+^ T cells in the BAL of unvaccinated RMs, suggesting migration to the primary infection site. This increase correlated with higher pathology scores compared to vaccinated RMs, hinting that HLA-E/*Mtb* CD3^+^ T cells might serve as a marker for *Mtb* infection. HLA-E CMV CD3^+^ T cells were detected in the BAL of each RM as well, which is surprising as CMV does not infect the airways like *Mtb*. HLA-E/*Mtb* CD4^+^ and CD8^+^ T cell frequencies remained unchanged in the circulation after *Mtb* challenge in BCG unvaccinated and vaccinated RMs, whereas in active TB and *Mtb* infected humans HLA-E/*Mtb* CD8^+^ T cell frequencies were higher relative to healthy controls, shown previously[13]. Disparities between species, *Mtb* strains and controlled exposure to a defined *Mtb* dose instead of naturally acquiring TB could account for the differences between humans and RMs. The route of administration for mucosal BCG vaccination and *Mtb* challenge in RMs was similar but only *Mtb* increased frequencies of HLA-E/*Mtb* CD4^+^ and CD8^+^ T cells, suggesting that differences in the infection cycle or virulence influenced the capacity to induce HLA-E/*Mtb* restricted T cells.

Our findings show that HLA-E CMV TMs were primarily recognized by CD8 expressing T cells, as described earlier[38,39], whereas HLA-E/*Mtb* TMs were recognized by both CD4 and CD8 expressing T cells that significantly correlated in both humans and RMs. Future studies should be directed to understand priming of HLA-E restricted T cells, as this likely deviates from conventional T cell priming, both for canonical peptides and especially for HLA-E restricted peptides that are sequentially unrelated to canonical peptides (pathogen-derived), also to establish if co-receptor independent recognition is an HLA-E specific phenomenon.

Due to poor staining of the memory markers on BAL samples, the memory phenotype could only be evaluated in PBMCs. Whereas the memory phenotype of circulating HLA-E/*Mtb* CD3^+^ T cells remained unchanged after BCG vaccination in RMs and BCG infection in humans (Figure S6) and after *Mtb* challenge in RMs, HLA-E CMV CD8^+^ T cells had an effector memory and terminally differentiated phenotype in RMs, possibly because of prolonged exposure to CMV via circulation in the RM colonies, confirming findings of previous studies[38,40]. The differential CCR expression pattern on HLA-E/*Mtb*, HLA-E/CMV and total T cells might suggest that HLA-E/*Mtb* T cells are more Th2-like T cells and HLA-E/CMV T cells more Th1-like T cells, as described earlier in humans[17,40]. Functional responses, such as the secretion of cytokines, which was not assessed because of limited material availability, should be evaluated for further substantiation.

While the frequency of HLA-E/*Mtb* CD4^+^ and CD8^+^ T cells was similar between *Mtb*/SIV co-infected and *Mtb* only infected CMs, we previously demonstrated that aTB/HIV co-infected individuals had the highest frequency of HLA-E/*Mtb* CD8^+^ T cells compared to individuals with aTB and TBI[13]. These individuals were not recently vaccinated with BCG as in CMs and the order of infections as controlled in the CM study was unknown in humans, which could explain the difference between CMs and the previous results on human PBMC samples.

Prominent differences were observed between T cells recognizing HLA-E CMV and HLA-E/*Mtb* in terms of frequencies, memory phenotype and co-receptor expression, which precludes that the findings for HLA-E/*Mtb* T cells were merely an effect of non-specific binding or stickiness of the HLA-E/*Mtb* TMs. The frequencies of HLA-E/*Mtb* T cells were low and might have been increased by including more HLA-E/*Mtb* peptides than the four included in this study. It is known that TCR affinity for HLA-E/peptide complexes in general, but especially for peptides that are different to self-peptides, is much lower than for classical HLA-I/peptide complexes, which can also account for the overall low frequencies of HLA-E/*Mtb* T cells[10,14,21]. Furthermore, our results were limited by the number of animals and humans included per study, in particular in the *Mtb*/SIV co-infection study, and the limited material that was available per individual/NHP.

In contrast to classical HLA-I molecules, HLA-E is not downregulated by HIV, has limited genetic diversity, can bind to *Mtb* peptides and can be recognized by T cells. These advantages highlight HLA-E’s potential as a vaccine target, either to induce *de novo* responses in BCG vaccinated individuals or as a primary vaccine, for example, via vaccination with various immunogenic *Mtb* epitopes in a formulation capable of inducing protective HLA-E restricted T cell responses. Hansen et al. already showed that MHC-E CD8^+^ T cells were essential to protect against SIV in SIV-infected RMs, but future studies are needed to assess the contribution of HLA-E restricted T cell responses in TB to confirm its efficacy as a primary target[24,41]. Consequently, our study expands current knowledge on the induction of HLA-E restricted T cells after BCG administration and *Mtb* infection and provides new insights into the exploration of HLA-E as a potential protective immune mechanism for TB.

**Supplementary Materials:** Figure S1: Timelines of the human and NHP studies; Figure S2: Gating strategies to determine HLA-E*01:01 and *01:03 CD4^+^ and CD8^+^ T cell frequencies and their memory phenotype in PBMCs of the human BCG revaccination study (A) and in PBMCs and the BAL of the human BCG infection studies (B-C); Figure S3: Gating strategies to determine HLA-E*01:03 CD4^+^ and CD8^+^ T cell frequencies, chemokine receptor expression and memory phenotype in PBMCs (A) and the BAL (B) of NHP samples in RM studies 1 and 2 and in PBMCs from CMs; Figure S4: Representative density plots to determine HLA-E T cells; Figure S5: Data of HLA-E*01:03 T cells for *Mtb* pool 1; Figure S6: Memory phenotype analysis of HLA-E*01:03 *Mtb* CD3^+^ T cells in BCG revaccinated healthy volunteers.

## Author contributions

Conceptualization, S.J. and T.O.; methodology, L.V., M.W., A.G., I.S., J.M., K.D., A.W., S.S., H.S., K.M., T.S.; validation, S.J., T.O., F.W., T.S., S.S. and H.S.; formal analysis, L.V. and M.W.; investigation, L.V., M.W., A.G., I.S., J.M., F.W., K.D., A.W.; resources, A.G., I.S., J.M., F.W., K.F., K.D., A.W., S.S., H.S., T.S.; data curation, L.V., M.W., I.S., A.G., F.W., A.W., S.S.; writing—original draft preparation, L.V., S.J., K.M., M.W.; writing—review and editing, L.V., S.J., K.M., M.W., T.O., K.F., A.G., T.S., I.S., J.M., K.D., F.W., A.W., S.S., H.S.; visualization, L.V. and M.W.; supervision, S.J., T.O., T.S., H.S., F.W., S.S.; project administration, S.J., T.O., T.S., H.S., S.S., F.W.; funding acquisition, S.J. and T.O.. All authors have read and agreed to the published version of the manuscript.

## Funding

This research was supported by the National Institute of Allergy and Infectious Diseases of the National Institutes of Health (R01AI141315) to Tom H.M. Ottenhoff and Simone A. Joosten.

### Institutional Review Board Statement

The human BCG revaccination study was approved by the Medicines Control Council (MCC) of South Africa (now called South African Health Products Regulatory Authority, SAHPRA), by the University Hospital Cleveland Medical Center Institutional Board and the Human Research Ethics Committee of the University of Cape Town (387/2008). The trial protocol of the first BCG controlled human infection study was approved by the Medicines and Healthcare products Regulatory Agency (EudraCT: 2015-004981-27) and the South-Central Oxford A Research Ethics Committee (REC) (15/SC/0716). The trial protocol of the second aerosol BCG infection study was approved by the Central Oxford A REC (18/SC/0307) and is registered in ClinicalTrials.gov (NCT03912207). The first RM study was approved by the UK Health Security Agency Animal Welfare and Ethical Review Body and was authorised under UK Home Office Licence P219D8D1A. The study protocol of the second RM study approved by the independent ethics committee Dier Experimenten Commissie (DEC) (761subB) and the institutional animal welfare body of the Biomedical Primate Research Center (BPRC). The CM study design and animal housing was approved by the Establishment Animal Welfare and Ethical Review Committee and authorised under UK Home Office Project License P219D8D1A.

### Informed Consent Statement

Informed consent was obtained from all individuals involved in the study.

### Data Availability Statement

The original contributions presented in the study are included in the article/supplementary material, further inquiries can be directed to the corresponding author.

## Acknowledgments

We thank our colleagues from the Flow Core Facility of the LUMC for their assistance and help. We further thank Dr. Hazel Morrison from Oxford University who helped in the generation of the BAL samples. Cartoons were created with BioRender.com.

## Disclaimer/Publisher’s Note

The statements, opinions and data contained in all publications are solely those of the individual author(s) and contributor(s) and not of MDPI and/or the editor(s). MDPI and/or the editor(s) disclaim responsibility for any in-jury to people or property resulting from any ideas, methods, instructions or products referred to in the content.

## Supplementary Materials

**Figure S1.**
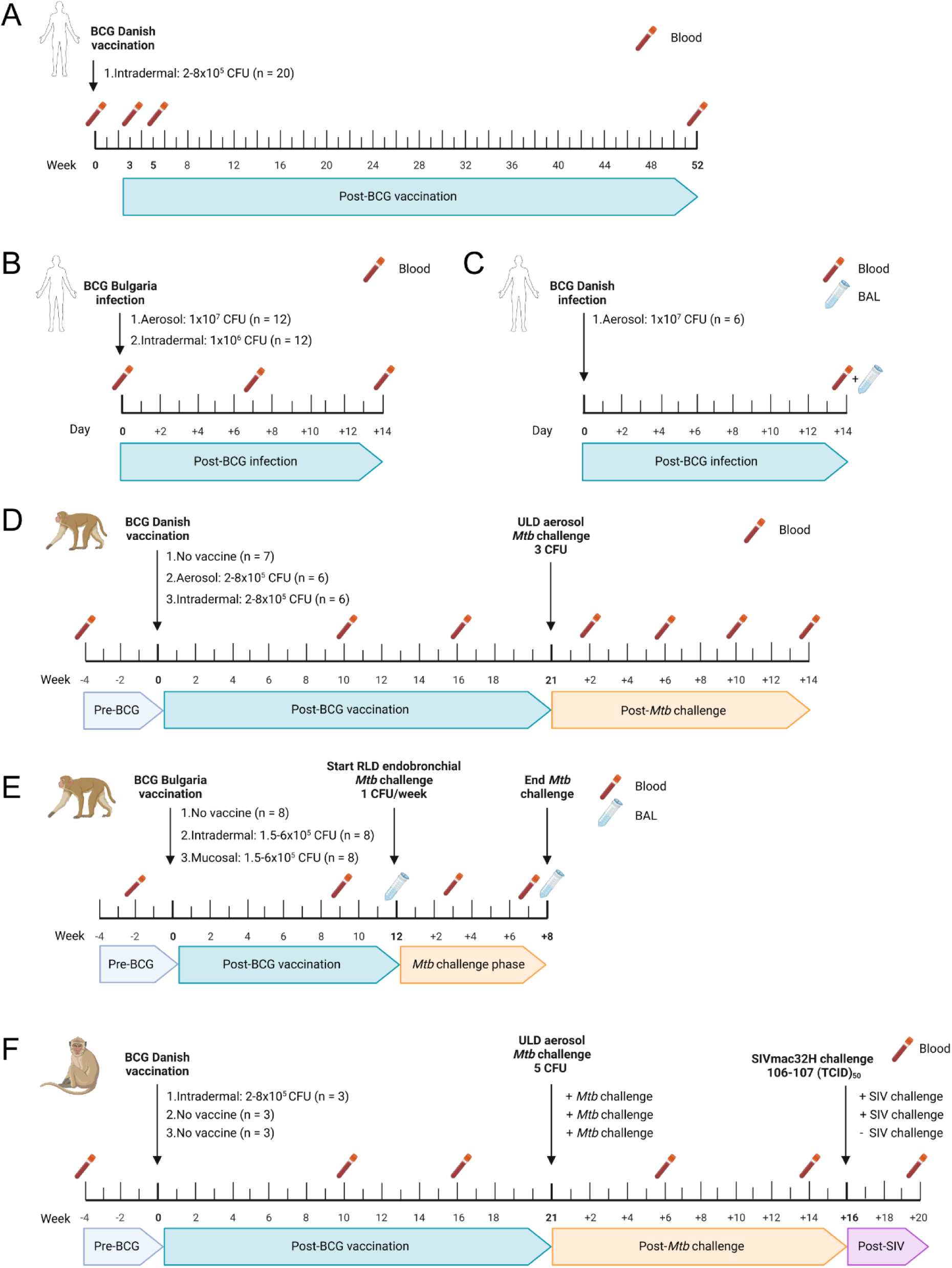
Timelines of the human and NHP studies. Timeline of the human BCG revaccination study. Blood was collected at indicated timepoints; **(A)** Timeline of the controlled human BCG infection study 1. Blood was collected at indicated timepoints; **(B)** Timeline of the controlled human BCG infection study 2. Blood and BAL were collected at indicated timepoints; **(C)** Timeline of BCG vaccination and *Mtb* challenge in RM study 1. Blood was collected at indicated timepoints; **(D)** Timeline of BCG vaccination and *Mtb* challenge in RM study 2. Blood and BAL were collected at indicated timepoints; **(E)** Timeline of BCG vaccination, *Mtb* challenge and SIV infection in the CM study. Blood was collected at indicated timepoints.

**Figure S2.**
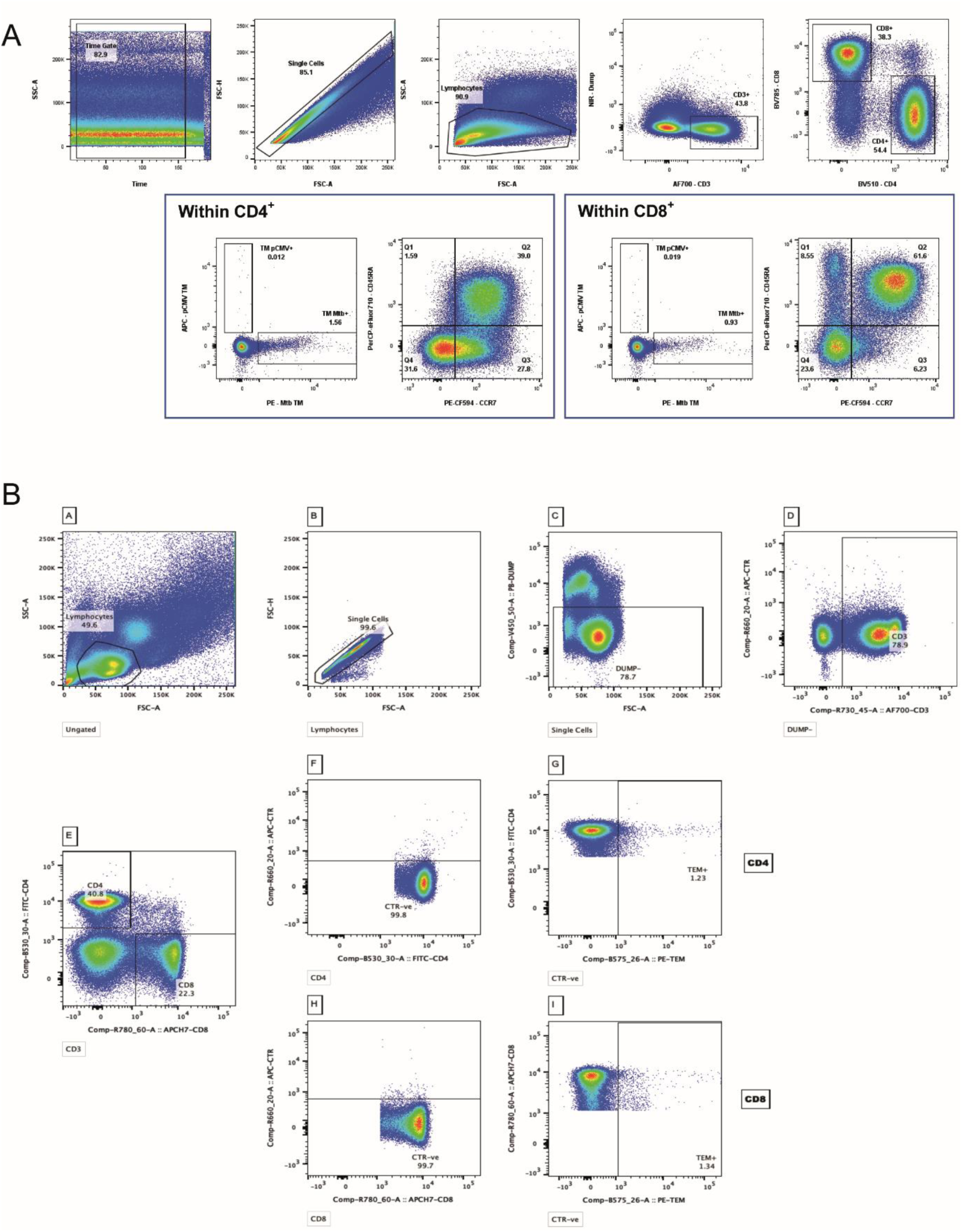
Gating strategies to determine HLA-E*01:01 and *01:03 CD4^+^ and CD8^+^ T cell frequencies and their memory phenotype in PBMCs of the human BCG revaccination study (A) and in PBMCs and BAL of the human BCG infection studies (B).

**Figure S3.**
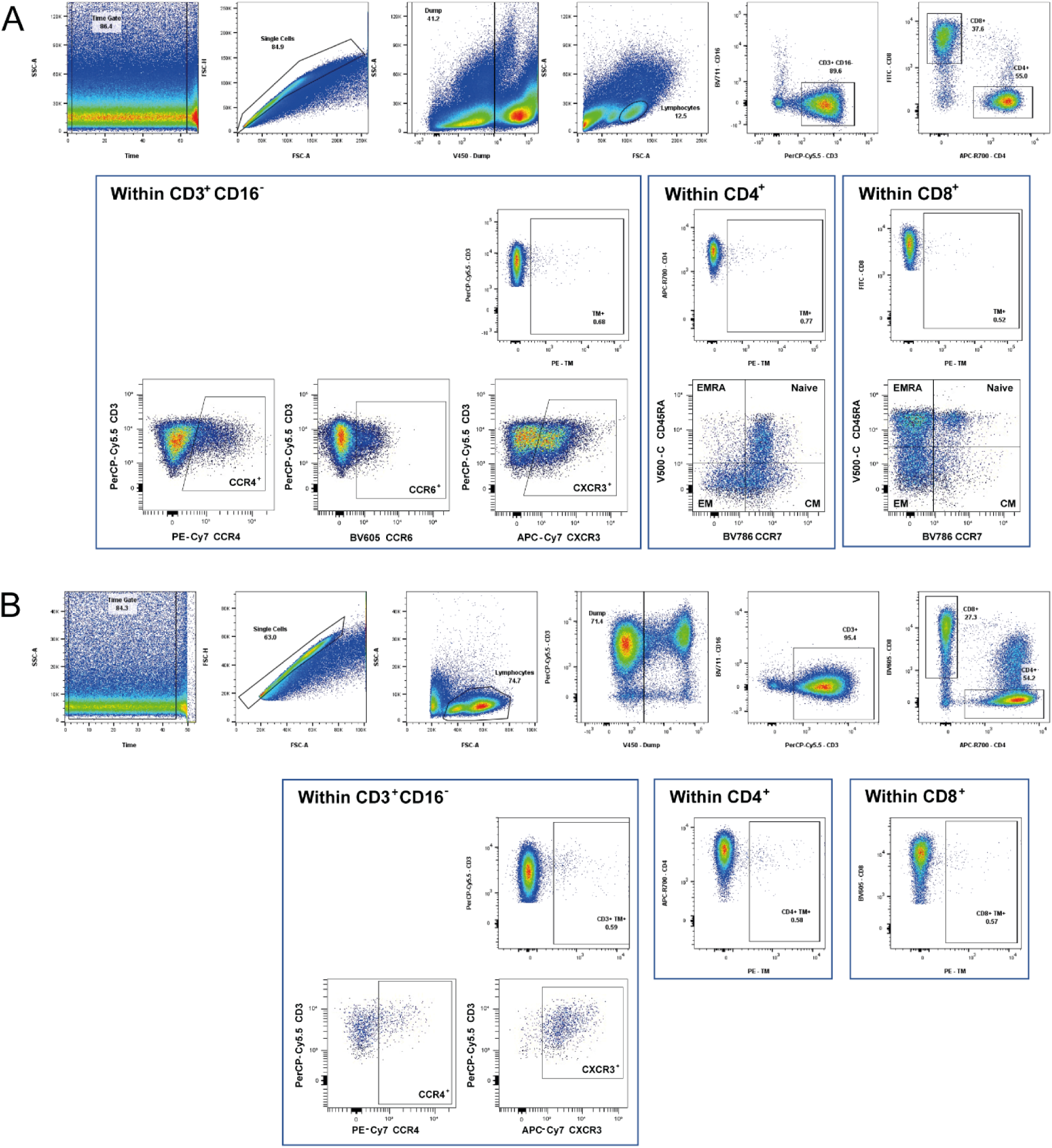
Gating strategies to determine HLA-E*01:03 CD4^+^ and CD8^+^ T cell frequencies, chemokine receptor expression and memory phenotype in PBMCs (A) and BAL (B) of NHP samples in RM studies 1 and 2 and in PBMCs from CMs.

**Figure S4.**
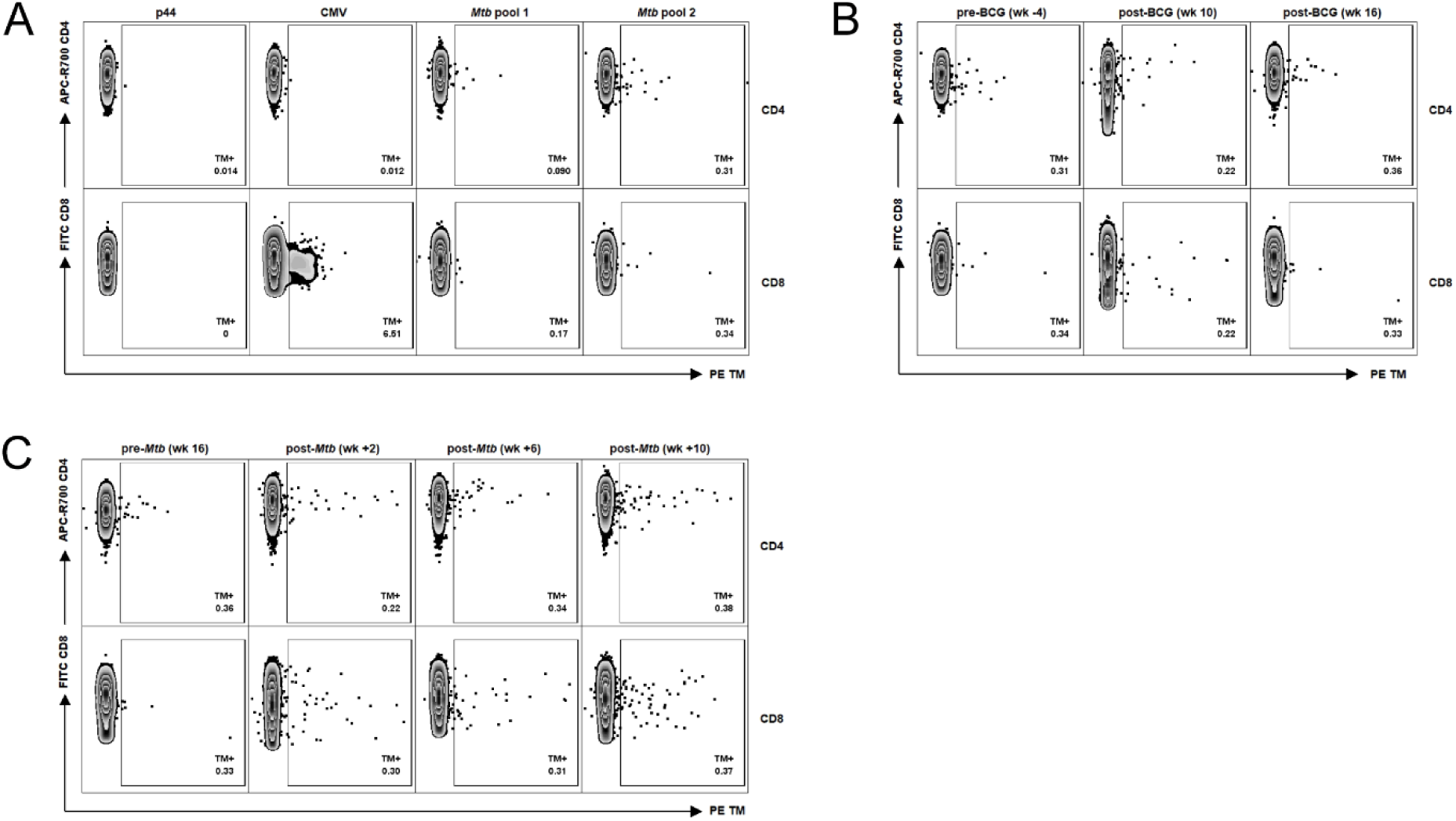
Representative density plots to determine HLA-E T cells. Peptide pools for HLA-E TM staining on RM samples are shown in Table 1. **(A)** Representative density plots for p44, CMV and *Mtb* pool 1 and 2 HLA-E*01:03 CD4^+^ and CD8^+^ T cell frequencies in one aerosol BCG vaccinated RM of study 1 at the pre-vaccination time point; **(B)** Representative density plots for *Mtb* pool 2 HLA-E*01:03 CD4^+^ and CD8^+^ T cell frequencies at the pre-vaccination time point and 10 and 16 weeks post-BCG vaccination in one aerosol vaccinated RM of study 1; **(C)** Representative density plots for *Mtb* pool 2 HLA-E*01:03 CD4^+^ and CD8^+^ T cell frequencies, 16 weeks post-BCG vaccination and 2, 6 and 10 weeks post-*Mtb* challenge in one aerosol vaccinated RM of study 1. In A-C, HLA-E TMs labelled with PE are shown on the X-axis and CD4 and CD8 are shown on the Y-axis.

**Figure S5.**
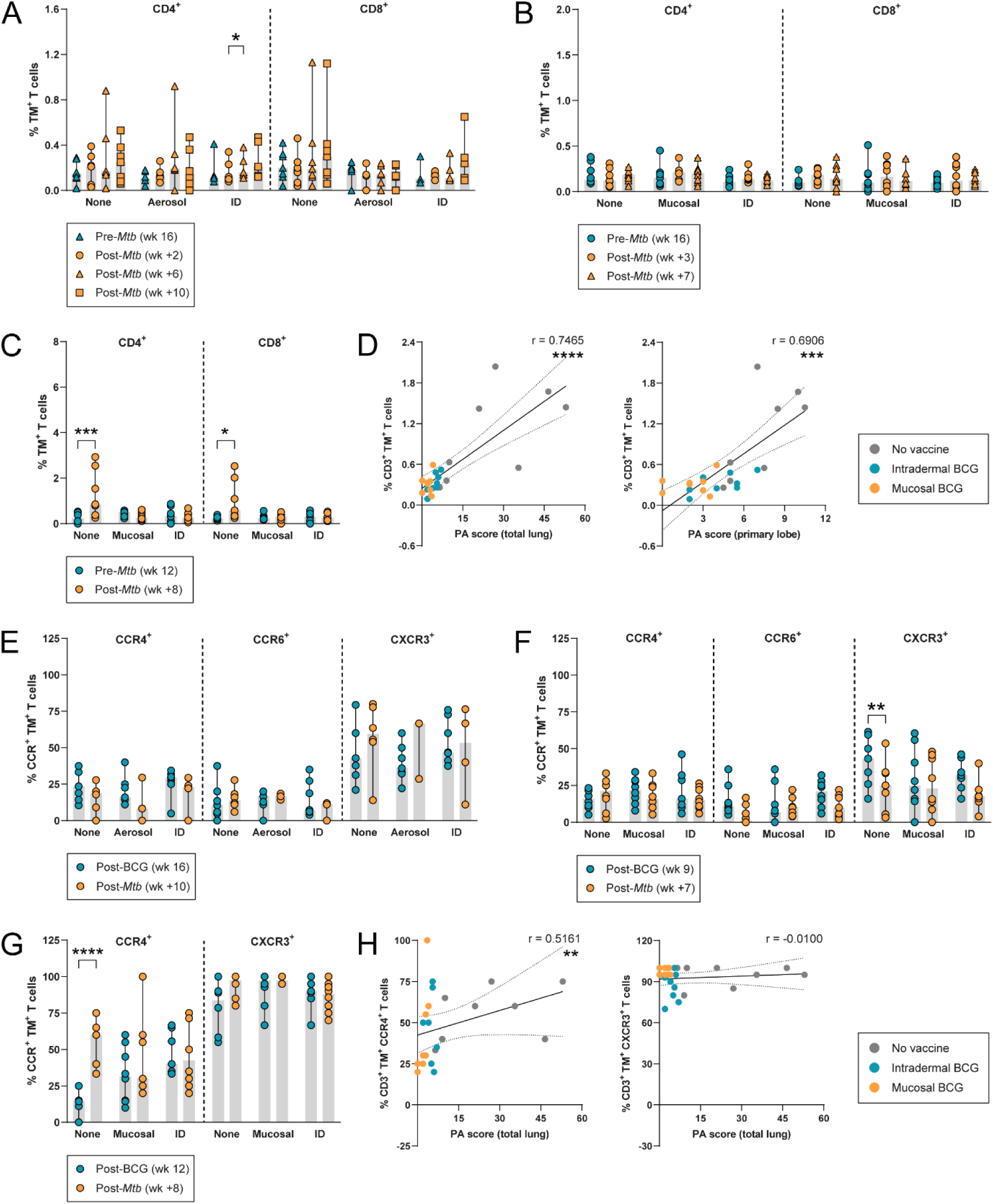
Data of HLA-E*01:03 T cells for *Mtb* pool 1. Peptides and sequences for HLA-E TM staining with *Mtb* pool 1 on RM samples are shown in Table 1. **(A)** HLA-E*01:03 CD4^+^ and CD8^+^ T cell frequencies in RM study 1 at the pre-*Mtb* challenge time point (blue triangles) and 2 (orange circles), 6 (orange triangles) and 10 weeks (orange squares) post-*Mtb* challenge; **(B)** Same as A, but then for RM study 2 at the pre-*Mtb* challenge time point (blue circles) and 3 and 7 weeks (orange circles and orange triangles) post-*Mtb* challenge; **(C)** HLA-E*01:03 CD4^+^ and CD8^+^ T cell frequencies in BAL of RM study 2 at the post-BCG vaccination time point (blue) and the post-*Mtb* challenge time point (orange); **(D)** Correlating the frequency of HLA-E*01:03 CD3^+^ T cells in BAL (Y-axis) and the PA scores in the total lung (left) and primary lobe (right) (X-axis) of RM study 2. Grey dots represent the unvaccinated group, blue dots the intradermal BCG vaccinated group and orange dots the mucosal BCG vaccinated group. Dotted lines represent the 95% confidence interval; **(E)** CCR4, CCR6 and CXCR3 expression on HLA-E*01:03 CD3^+^ T cells in RM study 1, 16 weeks post-BCG vaccination (blue) and 10 weeks post-*Mtb* challenge (orange); **(F)** Same as E, but then for RM study 2, 9 weeks post-BCG vaccination (blue) and 7 weeks post-*Mtb* challenge (orange); **(G)** Same as F (without CCR6), but then in BAL 12 weeks post-BCG vaccination (blue) and 8 weeks post-*Mtb* challenge (orange); **(H)** Correlating the frequency of CCR4^+^ (left) and CXCR3^+^ (right) HLA-E*01:03 CD3^+^ T cells in BAL on the Y-axis and the PA scores in the total lung on the X-axis, 8 weeks post-*Mtb* challenge in RM study 2. Grey dots represent the unvaccinated group, blue dots the intradermal BCG vaccinated group and orange dots the mucosal BCG vaccinated group. Dotted lines represent the 95% confidence interval. Shaded bars represent the median frequency and the error bars represent the 95% confidence interval. Significance was tested using a repeated measures (RM) two-way ANOVA with multiple comparison correction (A – C, E – G) and a Spearman’s rank correlation (D, H). * = *p*<0.05, ** = *p*<0.01 *** = *p*<0.001, **** = *p*<0.0001.

**Figure S6.**
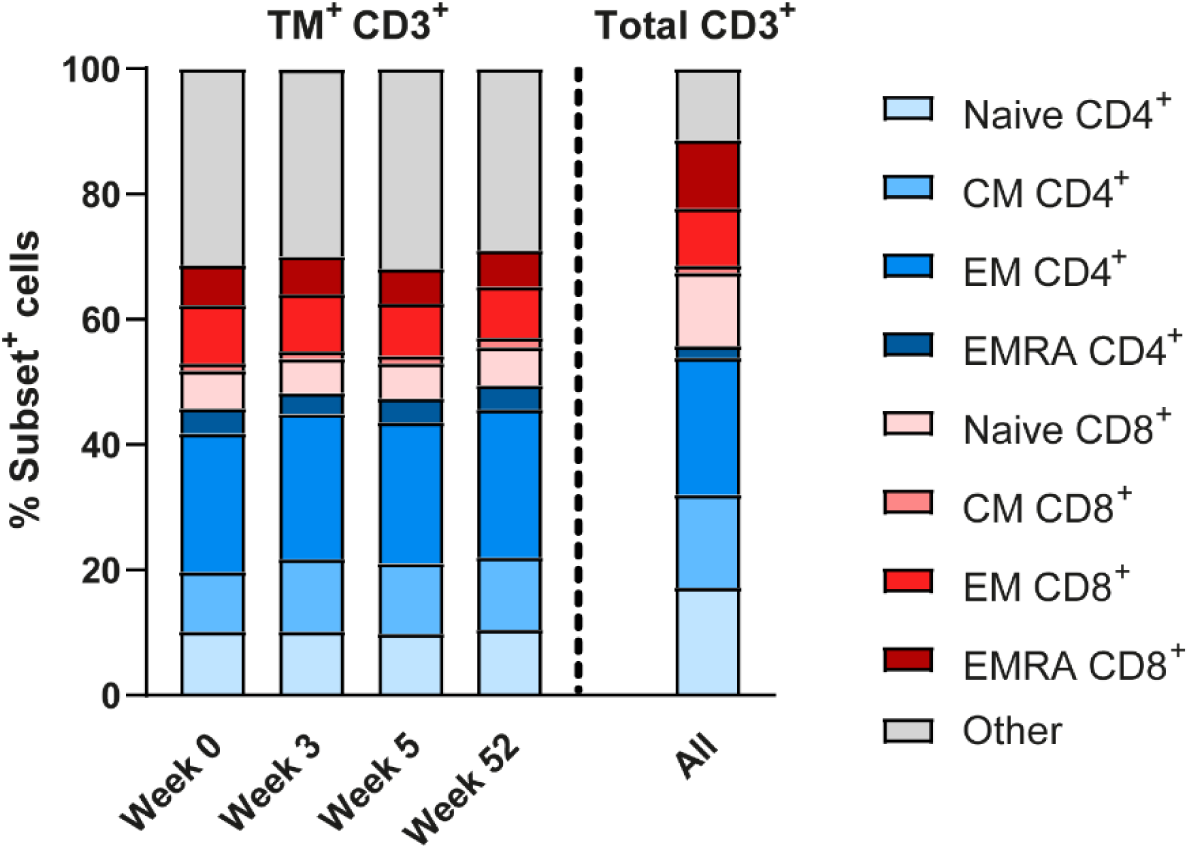
Memory phenotype analysis of HLA-E*01:03 *Mtb* CD3^+^ T cells in BCG revaccinated healthy volunteers. Peptides and sequences for HLA-E TM staining with the *Mtb* pool on human samples are shown in Table 1. Memory subset identification of HLA-E*01:03 CD3^+^ T cells for the *Mtb* pool relative to total CD3^+^ T cells in BCG revaccinated human volunteers (n=20) 0, 3, 5 and 52 weeks post-BCG revaccination relative to total CD3^+^ T cells. Stacked bars show the mean frequency.

## References

1. Global tuberculosis report 2023. Geneva: World Health Organization 2023.

2. Hunter, R.L.; Actor, J.K.; Hwang, S.A.; Khan, A.; Urbanowski, M.E.; Kaushal, D.; Jagannath, C. Pathogenesis and Animal Models of Post-Primary (Bronchogenic) Tuberculosis, A Review. Pathogens 2018, 7, doi:10.3390/pathogens7010019.

3. Cardona, P.J.; Williams, A. Experimental animal modelling for TB vaccine development. Int J Infect Dis 2017, 56, 268–273, doi:10.1016/j.ijid.2017.01.030.

4. Ruibal, P.; Voogd, L.; Joosten, S.A.; Ottenhoff, T.H.M. The role of donor-unrestricted T-cells, innate lymphoid cells, and NK cells in anti-mycobacterial immunity. Immunol Rev 2021, 301, 30–47, doi:10.1111/imr.12948.

5. Kanevskiy, L.; Erokhina, S.; Kobyzeva, P.; Streltsova, M.; Sapozhnikov, A.; Kovalenko, E. Dimorphism of HLA-E and its Disease Association. Int J Mol Sci 2019, 20, doi:10.3390/ijms20215496.

6. Strong, R.K.; Holmes, M.A.; Li, P.; Braun, L.; Lee, N.; Geraghty, D.E. HLA-E allelic variants. Correlating differential expression, peptide affinities, crystal structures, and thermal stabilities. J Biol Chem 2003, 278, 5082–5090, doi:10.1074/jbc.M208268200.

7. Lee, N.; Llano, M.; Carretero, M.; Ishitani, A.; Navarro, F.; Lopez-Botet, M.; Geraghty, D.E. HLA-E is a major ligand for the natural killer inhibitory receptor CD94/NKG2A. Proc Natl Acad Sci U S A 1998, 95, 5199–5204, doi:10.1073/pnas.95.9.5199.

8. Braud, V.M.; Allan, D.S.; O’Callaghan, C.A.; Soderstrom, K.; D’Andrea, A.; Ogg, G.S.; Lazetic, S.; Young, N.T.; Bell, J.I.; Phillips, J.H.; et al. HLA-E binds to natural killer cell receptors CD94/NKG2A, B and C. Nature 1998, 391, 795–799, doi:10.1038/35869.

9. Voogd, L.; Ruibal, P.; Ottenhoff, T.H.M.; Joosten, S.A. Antigen presentation by MHC-E: a putative target for vaccination? Trends Immunol 2022, 43, 355–365, doi:10.1016/j.it.2022.03.002.

10. Voogd, L.; Drittij, A.; Dingenouts, C.K.E.; Franken, K.; Unen, V.V.; van Meijgaarden, K.E.; Ruibal, P.; Hagedoorn, R.S.; Leitner, J.A.; Steinberger, P.;, et al. Mtb HLA-E-tetramer-sorted CD8(+) T cells have a diverse TCR repertoire. iScience 2024, 27, 109233, doi:10.1016/j.isci.2024.109233.

11. Ruibal, P.; Franken, K.; van Meijgaarden, K.E.; van Wolfswinkel, M.; Derksen, I.; Scheeren, F.A.; Janssen, G.M.C.; van Veelen, P.A.; Sarfas, C.; White, A.D.;, et al. Identification of HLA-E Binding Mycobacterium tuberculosis-Derived Epitopes through Improved Prediction Models. J Immunol 2022, 209, 1555–1565, doi:10.4049/jimmunol.2200122.

12. Ruibal, P.; Derksen, I.; van Wolfswinkel, M.; Voogd, L.; Franken, K.; El Hebieshy, A.F.; van Hall, T.; Schoufour, T.A.W.; Wijdeven, R.H.; Ottenhoff, T.H.M.;, et al. Thermal-exchange HLA-E multimers reveal specificity in HLA-E and NKG2A/CD94 complex interactions. Immunology 2023, 168, 526–537, doi:10.1111/imm.13591.

13. Prezzemolo, T.; van Meijgaarden, K.E.; Franken, K.; Caccamo, N.; Dieli, F.; Ottenhoff, T.H.M.; Joosten, S.A. Detailed characterization of human Mycobacterium tuberculosis specific HLA-E restricted CD8(+) T cells. Eur J Immunol 2018, 48, 293–305, doi:10.1002/eji.201747184.

14. Yang, H.; Sun, H.; Brackenridge, S.; Zhuang, X.; Wing, P.A.C.; Quastel, M.; Walters, L.; Garner, L.; Wang, B.; Yao, X.;, et al. HLA-E-restricted SARS-CoV-2-specific T cells from convalescent COVID-19 patients suppress virus replication despite HLA class Ia down-regulation. Sci Immunol 2023, 8, eabl8881, doi:10.1126/sciimmunol.abl8881.

15. Salerno-Goncalves, R.; Fernandez-Vina, M.; Lewinsohn, D.M.; Sztein, M.B. Identification of a human HLA-E-restricted CD8+ T cell subset in volunteers immunized with Salmonella enterica serovar Typhi strain Ty21a typhoid vaccine. J Immunol 2004, 173, 5852–5862, doi:10.4049/jimmunol.173.9.5852.

16. Vietzen, H.; Furlano, P.L.; Cornelissen, J.J.; Bohmig, G.A.; Jaksch, P.; Puchhammer-Stockl, E. HLA-E-restricted immune responses are crucial for the control of EBV infections and the prevention of PTLD. Blood 2023, 141, 1560–1573, doi:10.1182/blood.2022017650.

17. van Meijgaarden, K.E.; Haks, M.C.; Caccamo, N.; Dieli, F.; Ottenhoff, T.H.; Joosten, S.A. Human CD8+ T-cells recognizing peptides from Mycobacterium tuberculosis (Mtb) presented by HLA-E have an unorthodox Th2-like, multifunctional, Mtb inhibitory phenotype and represent a novel human T-cell subset. PLoS Pathog 2015, 11, e1004671, doi:10.1371/journal.ppat.1004671.

18. Joosten, S.A.; van Meijgaarden, K.E.; van Weeren, P.C.; Kazi, F.; Geluk, A.; Savage, N.D.; Drijfhout, J.W.; Flower, D.R.; Hanekom, W.A.; Klein, M.R.;, et al. Mycobacterium tuberculosis peptides presented by HLA-E molecules are targets for human CD8 T-cells with cytotoxic as well as regulatory activity. PLoS Pathog 2010, 6, e1000782, doi:10.1371/journal.ppat.1000782.

19. Doorduijn, E.M.; Sluijter, M.; Marijt, K.A.; Querido, B.J.; van der Burg, S.H.; van Hall, T. T cells specific for a TAP-independent self-peptide remain naive in tumor-bearing mice and are fully exploitable for therapy. Oncoimmunology 2018, 7, e1382793, doi:10.1080/2162402X.2017.1382793.

20. Doorduijn, E.M.; Sluijter, M.; Querido, B.J.; Seidel, U.J.E.; Oliveira, C.C.; van der Burg, S.H.; van Hall, T. T Cells Engaging the Conserved MHC Class Ib Molecule Qa-1(b) with TAP-Independent Peptides Are Semi-Invariant Lymphocytes. Front Immunol 2018, 9, 60, doi:10.3389/fimmu.2018.00060.

21. Yang, H.; Rei, M.; Brackenridge, S.; Brenna, E.; Sun, H.; Abdulhaqq, S.; Liu, M.K.P.; Ma, W.; Kurupati, P.; Xu, X.;, et al. HLA-E-restricted, Gag-specific CD8(+) T cells can suppress HIV-1 infection, offering vaccine opportunities. Sci Immunol 2021, 6, doi:10.1126/sciimmunol.abg1703.

22. Jorgensen, P.B.; Livbjerg, A.H.; Hansen, H.J.; Petersen, T.; Hollsberg, P. Epstein-Barr virus peptide presented by HLA-E is predominantly recognized by CD8(bright) cells in multiple sclerosis patients. PLoS One 2012, 7, e46120, doi:10.1371/journal.pone.0046120.

23. Bian, Y.; Shang, S.; Siddiqui, S.; Zhao, J.; Joosten, S.A.; Ottenhoff, T.H.M.; Cantor, H.; Wang, C.R. MHC Ib molecule Qa-1 presents Mycobacterium tuberculosis peptide antigens to CD8+ T cells and contributes to protection against infection. PLoS Pathog 2017, 13, e1006384, doi:10.1371/journal.ppat.1006384.

24. Hansen, S.G.; Marshall, E.E.; Malouli, D.; Ventura, A.B.; Hughes, C.M.; Ainslie, E.; Ford, J.C.; Morrow, D.; Gilbride, R.M.; Bae, J.Y.;, et al. A live-attenuated RhCMV/SIV vaccine shows long-term efficacy against heterologous SIV challenge. Sci Transl Med 2019, 11, doi:10.1126/scitranslmed.aaw2607.

25. Malouli, D.; Gilbride, R.M.; Wu, H.L.; Hwang, J.M.; Maier, N.; Hughes, C.M.; Newhouse, D.; Morrow, D.; Ventura, A.B.; Law, L.;, et al. Cytomegalovirus-vaccine-induced unconventional T cell priming and control of SIV replication is conserved between primate species. Cell Host Microbe 2022, 30, 1207–1218 e1207, doi:10.1016/j.chom.2022.07.013.

26. Verweij, M.C.; Hansen, S.G.; Iyer, R.; John, N.; Malouli, D.; Morrow, D.; Scholz, I.; Womack, J.; Abdulhaqq, S.; Gilbride, R.M.;, et al. Modulation of MHC-E transport by viral decoy ligands is required for RhCMV/SIV vaccine efficacy. Science 2021, 372, doi:10.1126/science.abe9233.

27. Nattermann, J.; Nischalke, H.D.; Hofmeister, V.; Kupfer, B.; Ahlenstiel, G.; Feldmann, G.; Rockstroh, J.; Weiss, E.H.; Sauerbruch, T.; Spengler, U. HIV-1 infection leads to increased HLA-E expression resulting in impaired function of natural killer cells. Antivir Ther 2005, 10, 95–107, doi:10.1177/135965350501000107.

28. Ruibal, P.; Franken, K.; van Meijgaarden, K.E.; Walters, L.C.; McMichael, A.J.; Gillespie, G.M.; Joosten, S.A.; Ottenhoff, T.H.M. Discovery of HLA-E-Presented Epitopes: MHC-E/Peptide Binding and T-Cell Recognition. Methods Mol Biol 2022, 2574, 15–30, doi:10.1007/978-1-0716-2712-9_2.

29. Gela, A.; Murphy, M.; Rodo, M.; Hadley, K.; Hanekom, W.A.; Boom, W.H.; Johnson, J.L.; Hoft, D.F.; Joosten, S.A.; Ottenhoff, T.H.M.;, et al. Effects of BCG vaccination on donor unrestricted T cells in two prospective cohort studies. EBioMedicine 2022, 76, 103839, doi:10.1016/j.ebiom.2022.103839.

30. Hatherill, M.; Geldenhuys, H.; Pienaar, B.; Suliman, S.; Chheng, P.; Debanne, S.M.; Hoft, D.F.; Boom, W.H.; Hanekom, W.A.; Johnson, J.L. Safety and reactogenicity of BCG revaccination with isoniazid pretreatment in TST positive adults. Vaccine 2014, 32, 3982–3988, doi:10.1016/j.vaccine.2014.04.084.

31. Satti, I.; Marshall, J.L.; Harris, S.A.; Wittenberg, R.; Tanner, R.; Lopez Ramon, R.; Wilkie, M.; Ramos Lopez, F.; Riste, M.; Wright, D.;, et al. Safety of a controlled human infection model of tuberculosis with aerosolised, live-attenuated Mycobacterium bovis BCG versus intradermal BCG in BCG-naive adults in the UK: a dose-escalation, randomised, controlled, phase 1 trial. Lancet Infect Dis 2024, doi:10.1016/S1473-3099(24)00143-9.

32. White, A.D.; Sarfas, C.; Sibley, L.S.; Gullick, J.; Clark, S.; Rayner, E.; Gleeson, F.; Catala, M.; Nogueira, I.; Cardona, P.J.;, et al. Protective Efficacy of Inhaled BCG Vaccination Against Ultra-Low Dose Aerosol M. tuberculosis Challenge in Rhesus Macaques. Pharmaceutics 2020, 12, doi:10.3390/pharmaceutics12050394.

33. Dijkman, K.; Sombroek, C.C.; Vervenne, R.A.W.; Hofman, S.O.; Boot, C.; Remarque, E.J.; Kocken, C.H.M.; Ottenhoff, T.H.M.; Kondova, I.; Khayum, M.A.;, et al. Prevention of tuberculosis infection and disease by local BCG in repeatedly exposed rhesus macaques. Nat Med 2019, 25, 255–262, doi:10.1038/s41591-018-0319-9.

34. White, A.D.; Sibley, L.; Gullick, J.; Sarfas, C.; Clark, S.; Fagrouch, Z.; Verschoor, E.; Salguero, F.J.; Dennis, M.; Sharpe, S. TB and SIV Coinfection; a Model for Evaluating Vaccine Strategies against TB Reactivation in Asian Origin Cynomolgus Macaques: A Pilot Study Using BCG Vaccination. Vaccines (Basel) 2021, 9, doi:10.3390/vaccines9090945.

35. van Wolfswinkel, M.; van Meijgaarden, K.E.; Ottenhoff, T.H.M.; Niewold, P.; Joosten, S.A. Extensive flow cytometric immunophenotyping of human PBMC incorporating detection of chemokine receptors, cytokines and tetramers. Cytometry A 2023, 103, 600–610, doi:10.1002/cyto.a.24727.

36. Walters, L.C.; Harlos, K.; Brackenridge, S.; Rozbesky, D.; Barrett, J.R.; Jain, V.; Walter, T.S.; O’Callaghan, C.A.; Borrow, P.; Toebes, M.;, et al. Pathogen-derived HLA-E bound epitopes reveal broad primary anchor pocket tolerability and conformationally malleable peptide binding. Nat Commun 2018, 9, 3137, doi:10.1038/s41467-018-05459-z.

37. Darrah, P.A.; Zeppa, J.J.; Maiello, P.; Hackney, J.A.; Wadsworth, M.H., 2nd; Hughes, T.K.; Pokkali, S.; Swanson, P.A., 2nd; Grant, N.L.; Rodgers, M.A.; et al. Prevention of tuberculosis in macaques after intravenous BCG immunization. Nature 2020, 577, 95–102, doi:10.1038/s41586-019-1817-8.

38. Sullivan, L.C.; Westall, G.P.; Widjaja, J.M.; Mifsud, N.A.; Nguyen, T.H.; Meehan, A.C.; Kotsimbos, T.C.; Brooks, A.G. The Presence of HLA-E-Restricted, CMV-Specific CD8+ T Cells in the Blood of Lung Transplant Recipients Correlates with Chronic Allograft Rejection. PLoS One 2015, 10, e0135972, doi:10.1371/journal.pone.0135972.

39. Pietra, G.; Romagnani, C.; Mazzarino, P.; Falco, M.; Millo, E.; Moretta, A.; Moretta, L.; Mingari, M.C. HLA-E-restricted recognition of cytomegalovirus-derived peptides by human CD8+ cytolytic T lymphocytes. Proc Natl Acad Sci U S A 2003, 100, 10896–10901, doi:10.1073/pnas.1834449100.

40. Mazzarino, P.; Pietra, G.; Vacca, P.; Falco, M.; Colau, D.; Coulie, P.; Moretta, L.; Mingari, M.C. Identification of effector-memory CMV-specific T lymphocytes that kill CMV-infected target cells in an HLA-E-restricted fashion. Eur J Immunol 2005, 35, 3240–3247, doi:10.1002/eji.200535343.

41. Hansen, S.G.; Wu, H.L.; Burwitz, B.J.; Hughes, C.M.; Hammond, K.B.; Ventura, A.B.; Reed, J.S.; Gilbride, R.M.; Ainslie, E.; Morrow, D.W.;, et al. Broadly targeted CD8(+) T cell responses restricted by major histocompatibility complex E. Science 2016, 351, 714–720, doi:10.1126/science.aac9475.

