## Supplementary figures and images for "*Mtb* specific HLA-E restricted T cells are induced during *Mtb* infection but not after BCG administration in non-human primates and humans"

### NHP - Supp Figure 1.tif

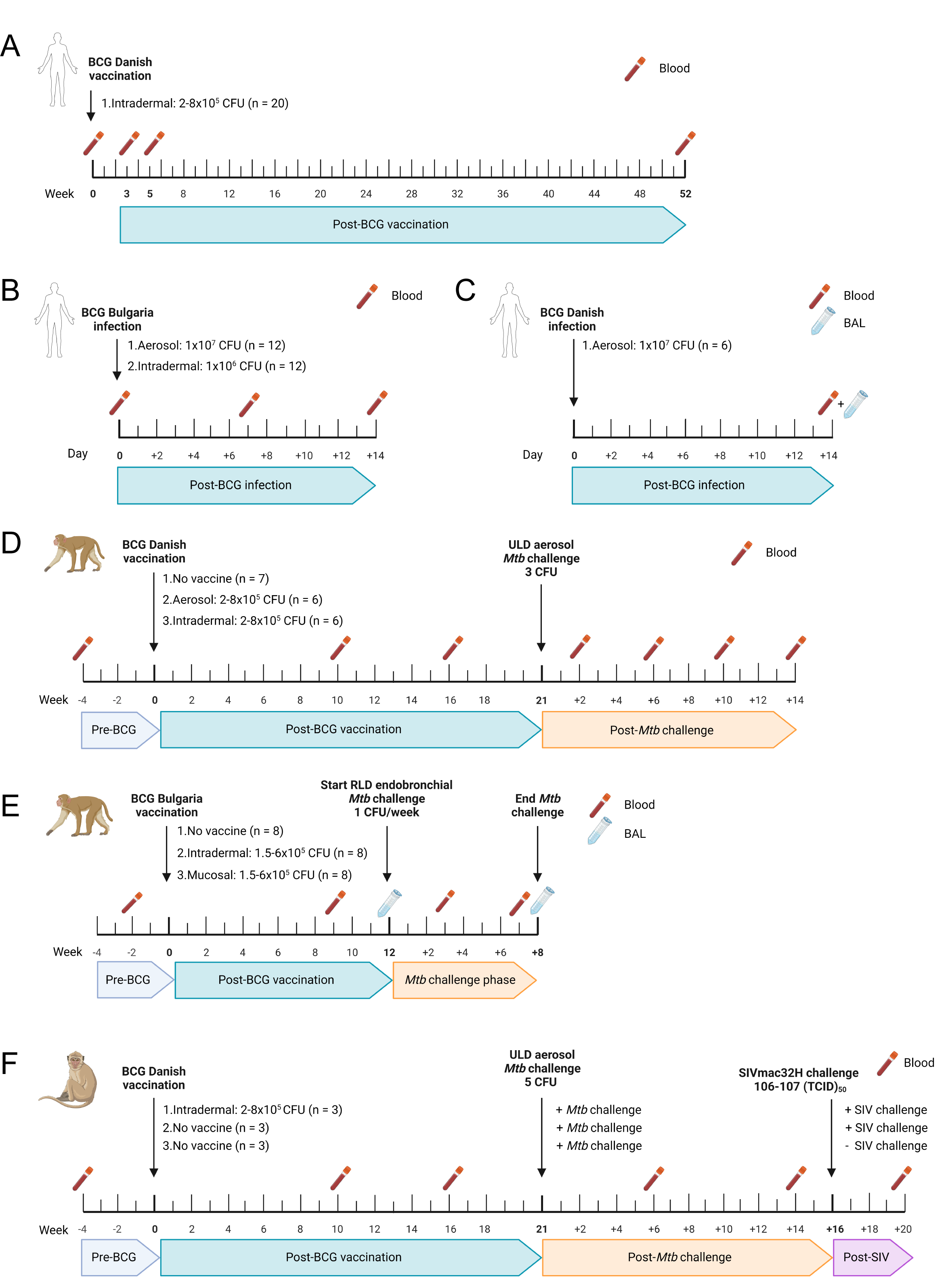

### NHP - Supp Figure 2.tif

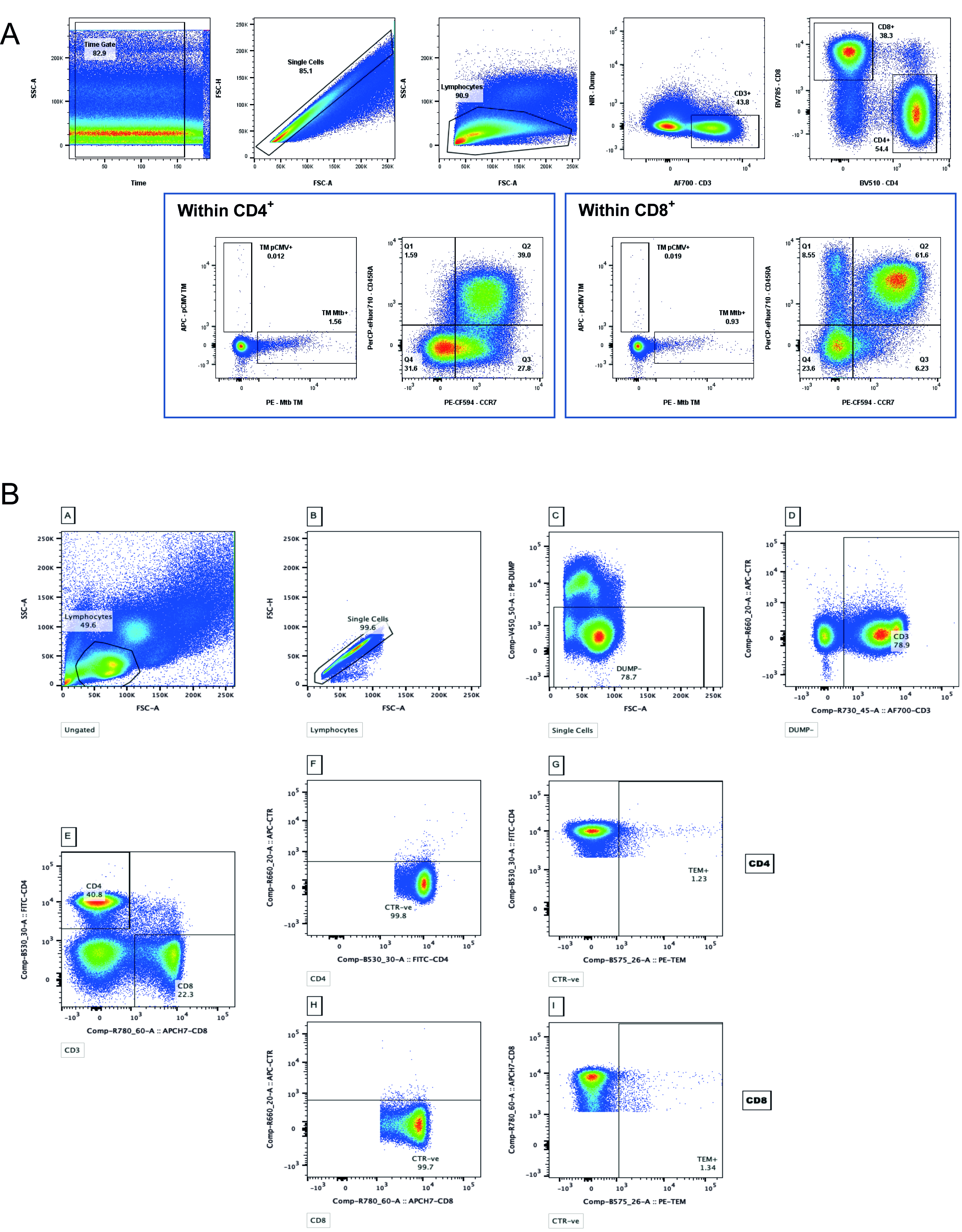

### NHP - Supp Figure 3.tif

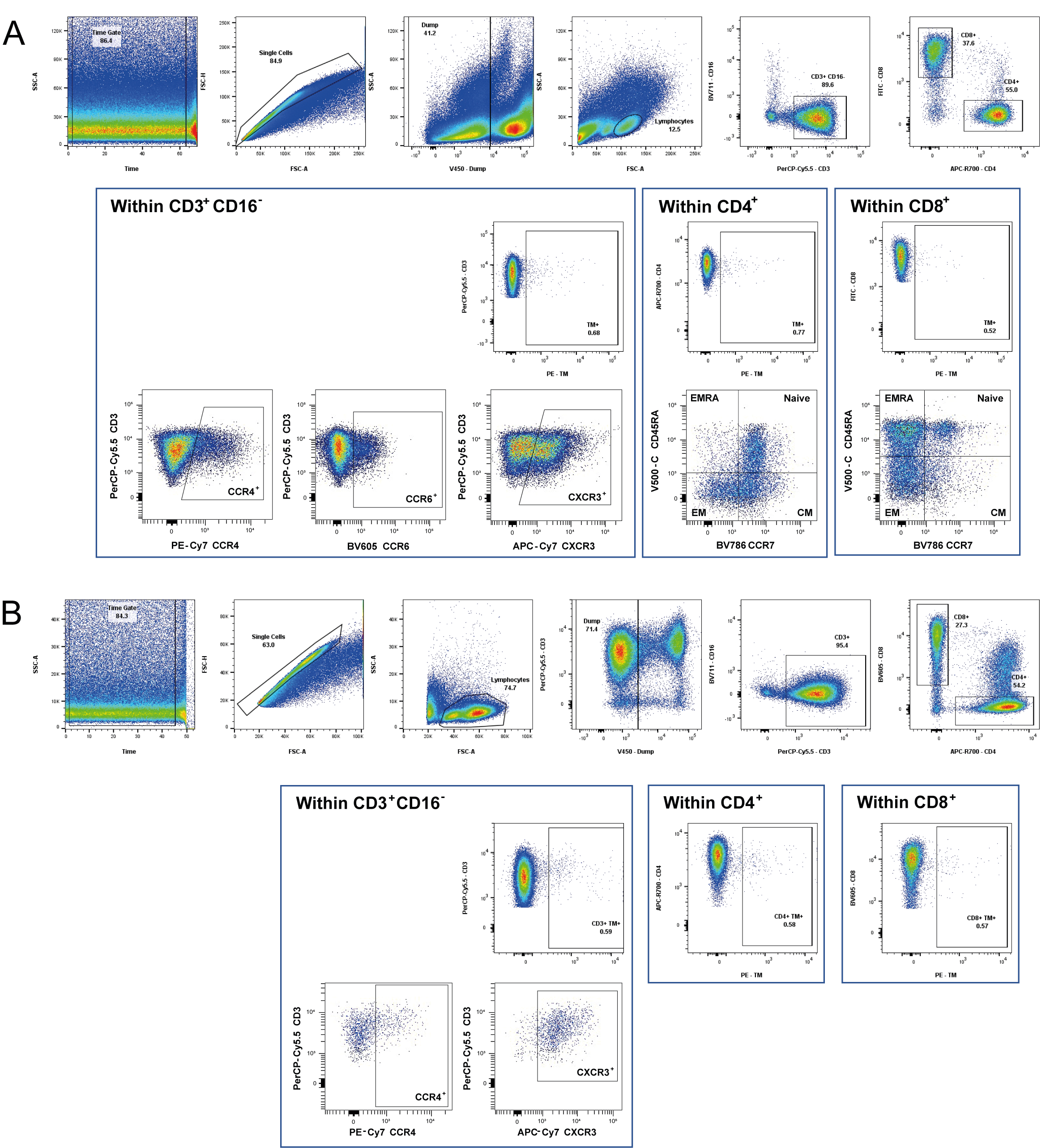

### NHP - Supp Figure 4.tif

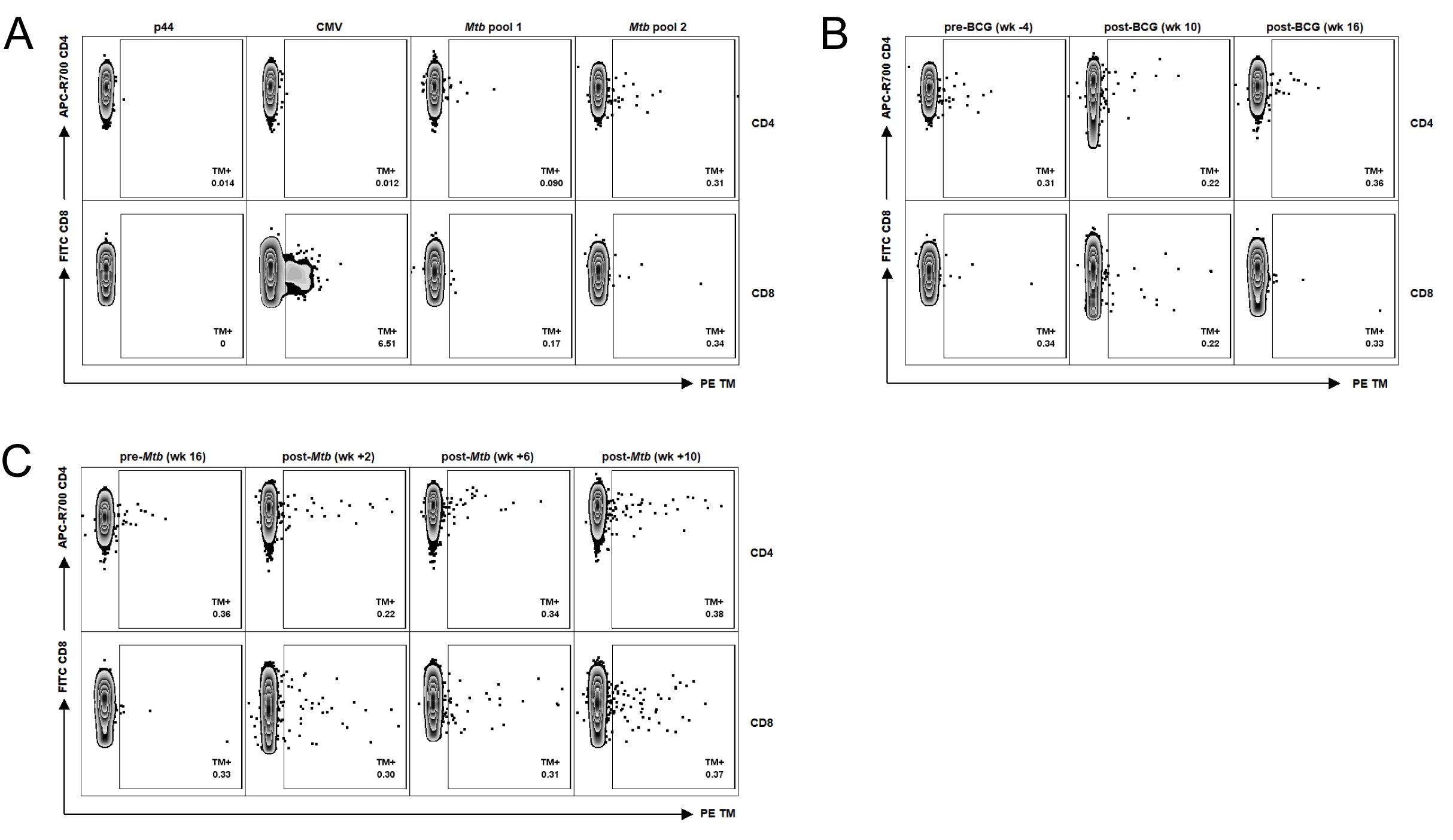

### NHP - Supp Figure 5.tif

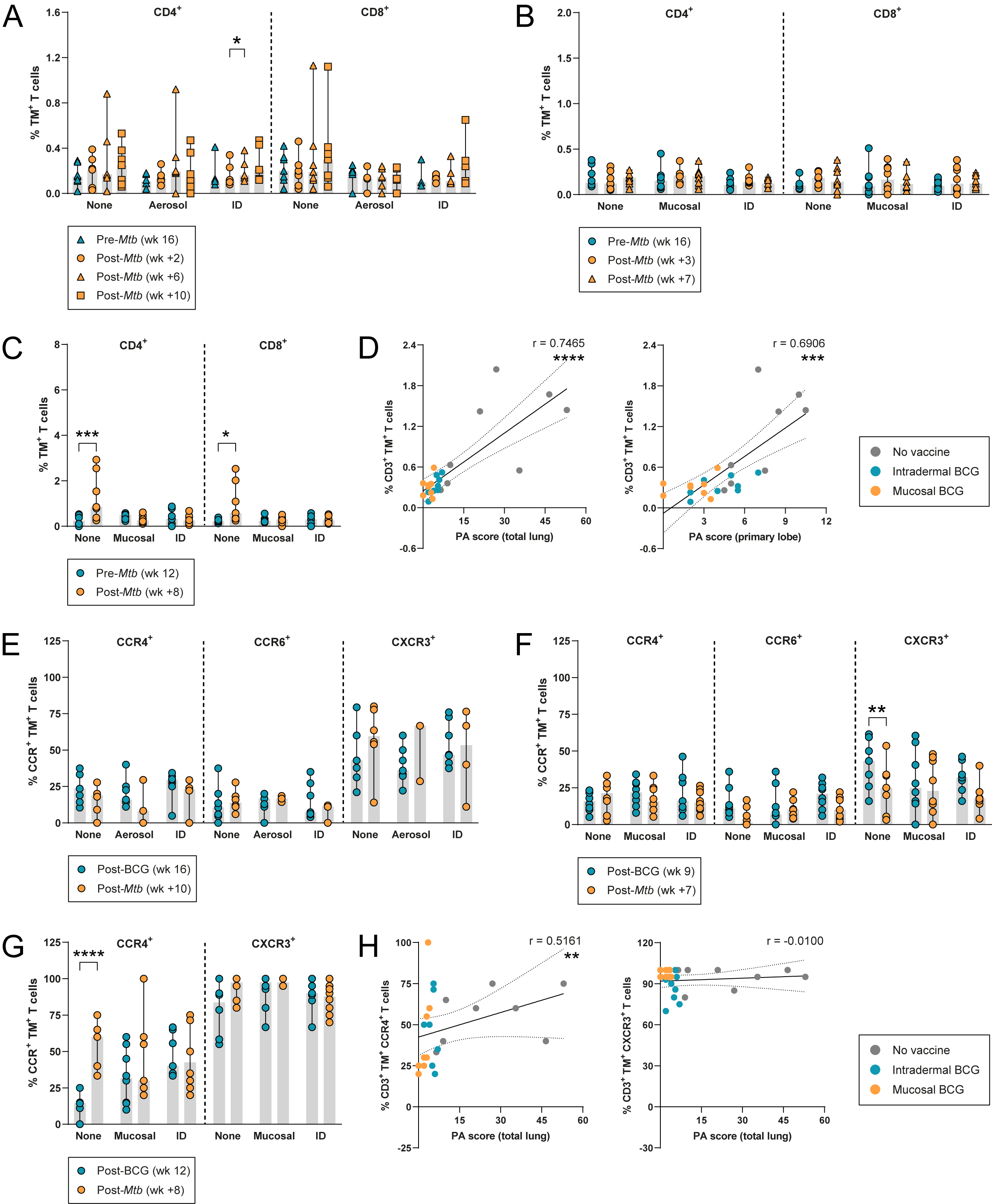

### NHP - Supp Figure 6.tif

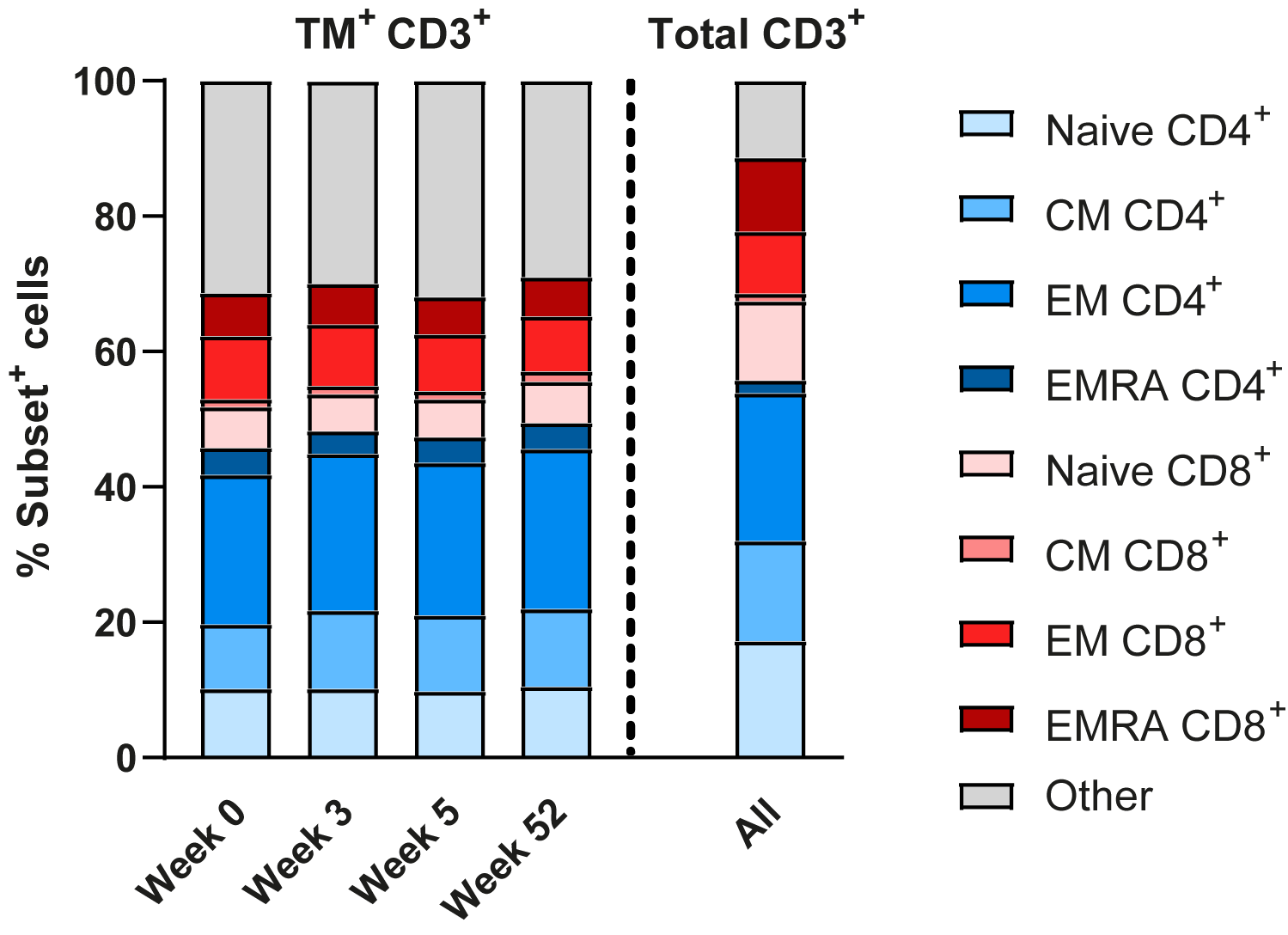
